# Gene drive and resilience through renewal with next generation *Cleave and Rescue* selfish genetic elements

**DOI:** 10.1101/2019.12.13.876169

**Authors:** Georg Oberhofer, Tobin Ivy, Bruce A Hay

## Abstract

Gene drive-based strategies for modifying populations face the problem that genes encoding cargo and the drive mechanism are subject to separation, mutational inactivation, and loss of efficacy. Resilience, an ability to respond to these eventualities in ways that restore population modification with functional genes is needed for long-term success. Here we show that resilience can be achieved through cycles of population modification with “*Cleave and Rescue”* (*ClvR*) selfish genetic elements. *ClvR* comprises a DNA sequence-modifying enzyme such as Cas9/gRNAs that disrupts endogenous versions of an essential gene, and a recoded version of the essential gene resistant to cleavage. *ClvR* spreads by creating conditions in which those lacking *ClvR* die because they lack functional versions of the essential gene. Cycles of modification can in principal be carried out if two *ClvR* elements targeting different essential genes are located at the same genomic position, and one of them, *ClvR*^n+1^, carries a *Rescue* transgene from an earlier element, *ClvR*^n^. *ClvR*^n+1^ should spread within a population of *ClvR*^n^, while also bringing about a decrease in its frequency. To test this hypothesis we first show that multiple *ClvR*s, each targeting a different essential gene, function when located at a common chromosomal position in *Drosophila*. We then show that when several of these also carry the *Rescue* from a different *ClvR*, they spread to transgene fixation in populations fixed for the latter, and at its expense. Therefore, genetic modifications of populations can be overwritten with new content, providing an ongoing point of control.

**Significance:** Gene drive can spread beneficial traits through populations, but will never be a one-shot project in which one genetic element provides all desired modifications, for an indefinitely long time. Here we show that gene drive mediated population modification in *Drosophila* can be overwritten with new content while eliminating old, using *Cleave and Rescue* (*ClvR*) selfish genetic elements. The ability to carry out cycles of modification that create and then leave behind a minimal genetic footprint while entering and exiting a population provides important points of control. It makes possible the replacement of broken elements, upgrades with new elements that better carry out their tasks and/or provide new functions, all while promoting the removal of modifications no longer needed.

## Introduction

Alleles of genes that confer desired traits are often unlikely to confer an overall fitness benefit on those that carry them (1, 2), particularly if the trait of interest ultimately results in death of carriers (3, 4). In consequence, specific strategies are needed to bring about an increase in the frequency of these genes in wild populations. Gene drive occurs when particular genetic elements – genes, gene complexes, or large chromosomal regions – are transmitted to viable, fertile progeny at rates greater than those of competing allelic variants or other parts of the genome. Transgenes, or alleles of endogenous loci, can be linked with a genetic element conferring drive, and this can promote their spread. A number of approaches to spreading traits through populations (population replacement/alteration/modification) in ways that are self-sustaining, by linking them with genetic elements that mediate drive, have been proposed (5–22). Several of these, *Medea* (*9, 23*), *UD^mel^* (15), engineered translocations (24), and *ClvR* (*Cleave and Rescue*) selfish genetic elements (25), have been implemented and shown to spread to transgene fixation in otherwise wildtype *Drosophila*. Sustained modification of a wildtype mosquito population using a homing based strategy, resulting in population suppression, has also been reported (26).

Any strategy to modify wild populations must contend with the inevitability of mutation and evolution in response to natural selection. Specifically, genes encoding cargo and constituting the drive mechanism are subject to separation and mutation to inactivity. Such mutations can result in loss of a functional cargo from the population if chromosomes carrying the inactive cargo, an empty drive element, or components of a drive element, are more fit than those carrying the full complement of active components. Evolution at other loci in the host, or in a pathogen the cargo is designed to target, can also occur such that the cargo becomes ineffective. Gene drive systems must be made robust – able to withstand forces leading to disruption – in order to delay the breakdown of a functional element. Examples of mechanisms to generate robustness in gene drive for population modification include multiplexing of components required for drive (25, 27, 28); interleaving of drive and Cargo components so as to prevent the creation of recombinant chromosomes that carry an empty drive element or an antidote only allele (in the case of drive elements that utilize a toxin and antidote) (9, 25); introducing multiple copies of a gene designed to inhibit pathogens, and by using multiple genes that target the pathogen through diverse mechanisms. However, these methods only delay failure, since none of them provides permanent protection against evolution through natural selection in response to all kinds of mutations and genetic diversity. Thus, population modification strategies must also be resilient – able to recover from a breakdown in ways that maintain or restore effective population modification over time. Releases into the wild of first generation elements may be dependent on the availability – or at least plausibility – of such strategies.

In principal resilience can be achieved if a new, second generation drive element can spread within a population fixed for an old element that has failed or lost efficacy. Because second generation elements are subject to the same evolutionary forces as first generation elements, it should also be possible to carry out additional cycles of modification. In consequence, a drive mechanism able to achieve resilience will likely use orthogonally acting components such that the presence of old drive elements does not interfere with drive by a newer generation element. For related reasons the components that make up a resilient drive mechanism should in some sense be indefinitely extensible in terms of the ability to create orthogonally acting new drive elements. Finally, an ideal system would not create unwanted genomic clutter from the accumulation of earlier generation non-functional elements and/or their components: cargo genes that have lost activity or have undesired effects; drive element components such as gRNAs that continue to create new LOF alleles at old essential gene loci, and which may compete for loading into Cas9 with gRNAs from a current generation element (see below); and dominant markers that serve no purpose and may interfere with monitoring the behavior of newer generation elements. Finally, a general cluttering of the genome with transgenes and mutant alleles is not compatible with what we consider to be a primary tenant of population modification – that it should be targeted, specific, and in some sense reversible, creating and then leaving behind a minimal genetic footprint as it enters and exits a population. In consequence, drive and population modification with a new element should result in a contemporaneous decrease in frequency of drive elements from earlier generations.

*ClvR* selfish genetic elements elements (25), also referred to as Toxin Antidote Recessive Embryo (TARE) in a related implementation (22), are good candidates for a system with these characteristics. They comprise two components, a DNA sequence-modifying enzyme such as Cas9 and gRNAs (the toxin/*Cleaver*) that acts in trans to disrupt the endogenous version of an essential gene through cleavage and inaccurate repair, and a recoded version of the essential gene resistant to cleavage (the antidote/*Rescue*). When these two components are tightly linked, Cas9 and gRNAs create potentially lethal loss of function (LOF) alleles of the essential gene, wherever it is located (Fig. 1A). However, the lethal LOF phenotype only manifests itself in those who fail to inherit *ClvR*. In contrast, those who inherit *ClvR* survive because they always inherit the recoded copy of the essential gene (Fig. 1B). Thus, *ClvR* spreads by killing those who lack it, creating a population that ultimately becomes dependent on – addicted to – the *ClvR*-encoded *Rescue* transgene. *ClvR* is in principal broadly extensible within a species since any gene that is essential for survival or fertility can be targeted for cleavage and rescue. These components are also orthogonally acting since Cas9/gRNAs only create mutations in the gene to which the gRNAs have homology, and the recoded *Rescue* transgene only rescues LOF phenotypes due to mutations induced by a gene-specific Cas9/gRNA complex.

**Fig 1.**
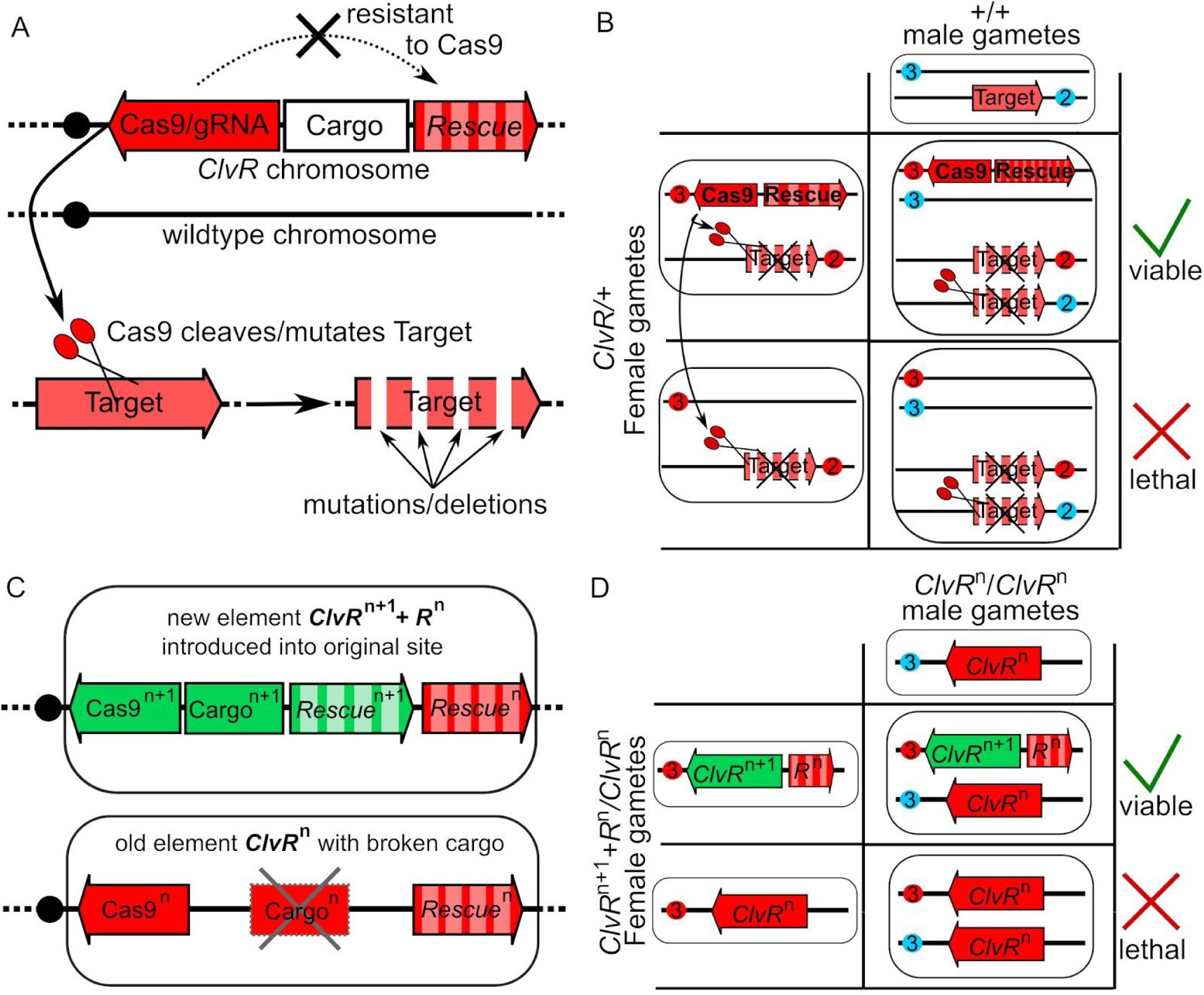
First and second generation *ClvR* elements and their genetic behavior. **(A)** Components that make up a *ClvR* element. See text for details. **(B)** A Punnett square highlighting the genetic mechanism by which *ClvR* elements bias inheritance in their favor. Maternal chromosomes are indicated with a red circle (centromere), paternal chromosomes in blue. In a heterozygous female, Cas9 (located in the *ClvR* on chromosome 3 in this example) cleaves and mutates to LOF the endogenous target gene on chromosome 2, in the germline. In addition, active Cas9/gRNA complexes are deposited maternally into all eggs. After mating with a wildtype male, maternally transmitted Cas9/gRNA cleaves/mutates the target gene on the paternal chromosome. Progeny that do not inherit *ClvR* and its recoded *Rescue* die because they lack essential gene function. **(C)** A female transheterozygous for *ClvR*^n^, which has an inactive Cargo^n^, and *ClvR*^n+1^, which carries a new Cargo^n+1^. **(D)** Gamete and progeny genotypes for a cross between the transheterozygous *ClvR*^n+1^/*ClvR*^n^ female and a homozygous *ClvR*^n^/*ClvR^n^* male. Chromosomes carrying the two different essential genes being targeted (as in panels A and B) are not indicated for clarity of presentation.

In earlier work we briefly outlined a general method by which cycles of population modification with the above characteristics – orthogonally acting components, indefinite extensibility, and contemporaneous removal of earlier generation elements – could in principle be carried out utilizing chromosomally located *Medea* (9, 29) or *ClvR* (25) selfish genetic elements, which each utilize a toxin-antidote based mechanism of action. In brief, these approaches involve creating a series of next generation drive elements in which each new element competes with, and ultimately displaces a first or earlier generation element. The basic strategy for carrying out cycles of modification with *ClvR* is illustrated in Fig. 1C, D. In this scenario the original *ClvR* is known as *ClvR*^n^. A second generation *ClvR*, *ClvR*^n+1^, which is meant to spread and supplant *ClvR^n^*, is located at the same position in the genome. Therefore, meiotic recombination cannot bring both *ClvR*s onto the same chromosome and they are forced to compete with each other for inheritance in viable progeny through the use of different combinations of Cas9/gRNA toxins and recoded *Rescue* antidotes. *ClvR^n^* targets essential gene^n^ for cleavage and rescue. *ClvR*^n+1^ targets essential gene^n+1^ for cleavage and rescue, while also carrying the *Rescue* transgene for essential gene^n^. Because progeny carrying *ClvR*^n^ or no *ClvR* element (wildtype) are sensitive to loss of essential gene^n+1^, those carrying *ClvR*^n+1^ have a survival advantage, regardless of their status with respect to *ClvR*^n^. Meiosis remains fair and both elements are inherited by progeny in normal Mendelian ratios, but the survival of progeny is biased towards those carrying *ClvR*^n+1^. As a result – and provided that the fitness costs associated with carrying *ClvR* are less than those experienced by non *ClvR*-bearing chromosomes in response to *ClvR*-dependent killing (25, 30) – the non-*ClvR*-bearing chromosome *ClvR*^n+1^ is expected to spread into populations of *ClvR*^n^ (and wildtype), while bringing about a corresponding decrease in the frequency of *ClvR*^n^. A Punnett square example that illustrates this behavior, in which a female transheterozygous for both elements mates with a *ClvR^n^* male, is presented in Fig. 1D.

Here we show that cycles of modification with *ClvR* elements can be achieved in *Drosophila*. We first show that when multiple *ClvR*s, each targeting a different essential gene, are located at a common chromosomal position, they show drive, resulting in rapid spread to transgene fixation in wildtype populations. We then show that when several of these *ClvR* elements elements also carry the *Rescue* transgene from a different element, a *ClvR^n^* element that has failed in some way and needs to be supplanted, the former – now a *ClvR^n+1^* element – spreads to transgene fixation at the expense of *ClvR^n^*, and at the expense of wildtype. These results show that *ClvR* is extensible with orthogonally acting components, and that population modifications can be overwritten with new instructions while eliminating old ones. These features provide important points of control with respect to replacement of broken elements, upgrades with new elements that better carry out their original jobs and/or provide new functions, and removal of old elements whose presence is no longer desired.

## Results

### Synthesis of two new *ClvR* elements at the same genomic position as *ClvR^tko^*

Cycles of population modification with *ClvR* elements can be carried out in two ways. First, *ClvR* elements could simply be introduced at new sites in the genome. If new elements freely recombine with old elements, and use orthogonally acting components (different gRNAs and essential genes), a new round of replacement should ensue in populations that carry one or more versions of earlier generation elements, at the same rate as for the first generation element. However, because the components of each *ClvR* are orthogonally acting, the earlier generation elements and their remnants will remain in the population at frequencies determined by natural selection. This creates the unwanted genomic clutter discussed above. Here we focus on the alternative strategy in which orthogonally acting *ClvR* elements are located at the same position in the genome, an arrangement that forces them to compete for survival.

This latter strategy demands that it be possible to create multiple, independently acting *ClvR* elements that show gene drive when located at a common position in the genome. We previously reported the creation of a single *ClvR* element, *ClvR^tko^*, in *D. melanogaster* (25). *ClvR^tko^* is located on the *D. melanogaster* third chromosome at map position 68E, spreads rapidly into wildtype *D. melanogaster* populations, and is functional in populations from five continents (25). To determine if new, orthogonally acting *ClvR*s can be created at this same genomic position, we synthesized two new *ClvR* elements using the same approach as for *ClvR^tko^*. The targeted essential genes were *dribble* (*dbe*) and *Transcription-factor-IIA-S* (*TfIIA-S*). *Dbe* is located at 21E2 on chromosome 2 and encodes a protein required for processing of cytoplasmic pre-ribosomal RNA (31). *TfIIA-S* is located at 95C8 on chromosome 3 and encodes a small subunit of a basal transcription factor that is a part of the *Pol-II* transcription machinery (32). Both genes are recessive lethal and expressed ubiquitously in *Drosophila*. For the *Rescue* component of *tko* we used the *tko* ortholog from the distantly related species *D. virilis*. For the new *ClvR* elements we used the target gene orthologs from *D. suzukii*, an agricultural pest of major economic importance. As with *ClvR^tko^*, the toxin/*Cleaver* part of the constructs consisted of Cas9 under the control of the germline specific *nanos* promoter, and 5’ and 3’ untranslated regions, and a set of four gRNAs designed to have homology with the *D.melanogaster* essential gene, but not the antidote/*Rescue* ortholog from *D. suzukii*, each expressed from a U6 promoter. The new *ClvR* elements also carried two dominant markers, ubiquitous *opie-td-tomato* and eye specific *3xP3-GFP*. A detailed description of construct assembly and fly germline transformation is given in the methods section and Fig. S1,2.

**Fig. 2.**
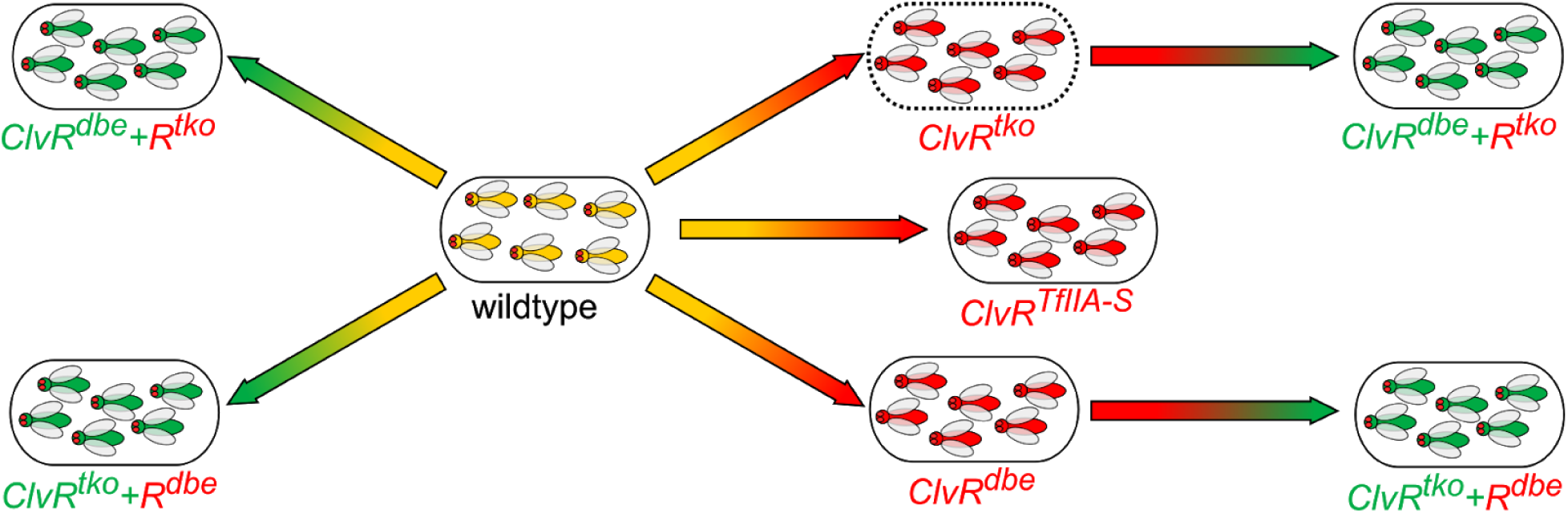
Gene drive experiments. Arrows proceed from wildtype, *ClvR^tko^*, or *ClvR^dbe^* to some other *ClvR*-bearing state. The blunt end of the arrow indicates the starting state and the pointed end the population state (transgene fixation) drive is meant to achieve. Color gradients schematically reflect progress from starting state to end state. Controls for each drive experiment are discussed in the text, and in Fig. 3,4. Experiments demonstrating drive of *ClvR^tko^* into a wildtype population (dashed outline) were published previously (25). See also Fig. S6 for further characterization of the *ClvR^tko^* experiment over more generations.

### Synthesis of 2nd Generation *ClvR^n+1^* elements

A second generation *ClvR*, *ClvR^n+1^*, consists of a *ClvR* that utilizes drive components that function orthogonally to those of *ClvR^n^* – a different toxin/*Cleaver* and a different antidote/*Rescue* – in addition to the antidote/*Rescue* from *ClvR^n^*. The new toxin/*Cleaver* and both antidote/*Rescues* must work well in order for such an element to spread in populations fixed for *ClvR^n^*. To create such elements we took flies that carried *ClvR^dbe^* and inserted into them the *Rescue* from *ClvR^tko^*, thereby creating a second generation element, *ClvR^dbe^+R^tko^*, designed to spread into populations of *ClvR^tko^* and wildtype. We also created the converse second generation *ClvR*, *ClvR^tko^+R^dbe^*, designed to spread into populations of *ClvR^dbe^* and wildtype. In each case we first assembled a construct that had the desired *Rescue^n^* fragment and a new dominant marker consisting of the ubiquitous *opie* promoter followed by a partial *GFP* open reading frame (ORF). These elements were flanked by homology arms matching the region surrounding the *3xP3* promoter driven *GFP* marker within the first generation *ClvR* element whose components would become Cas9/gRNA^n+1^ and *Rescue*^n+1^. First generation *ClvR* flies were injected with this donor plasmid along with Cas9 protein pre-loaded with a gRNA that binds between the *3xP3* promoter and the *GFP* open reading frame. Once Cas9 creates a DSB break between the *3xP3* promoter and *GFP*, the donor template can be used for repair, thereby, inserting the new *Rescue*. Positive transformants were identified by the change in *GFP* expression from eye-specific to ubiquitous. Correct insertion of *Rescue^n^* was confirmed by sequencing (see Fig. S1,2 for details).

### Genetic behavior of new 1st and 2nd Generation *ClvR* elements

We began our characterization of these four new *ClvR* elements by determining the frequency with which the target essential gene was mutated to LOF in crosses involving female and male *ClvR*-bearing parents. For females we crossed heterozygous *ClvR/+* (+ for wildtype locus) virgins to wildtype males (see the cross with generic elements depicted in Fig. 1B) and scored progeny for the presence of the dominant *ClvR* marker *td-tomato*. *ClvR* frequency was calculated as the number of *ClvR*-positive progeny divided by the total number of progeny. The cleavage rate to LOF is the fraction of *ClvR*-positive progeny divided by half the total number of progeny, since with mendelian inheritance 50% of the progeny would be expected to inherit *ClvR* in the absence of *ClvR*-dependent killing. These percentages are a function of maternal germline cleavage and cleavage in the embryo due to maternal carryover of Cas9/gRNAs. Males carrying *ClvR* do not show paternal carryover of Cas9 at appreciable frequencies (25). To reveal the LOF mutation status of target loci that are exposed to Cas9/gRNAs in the adult male germline we crossed heterozygous *ClvR/+* males to females that carried a deficiency (*Df*) for the target gene in trans to a balancer chromosome which is wildtype at the target locus. This allowed us to calculate the male germline LOF mutation creation rate directly in progeny as the percent of progeny carrying the *Df* for the essential gene that also carry *ClvR*, since those *Df*-bearing individuals not carrying *ClvR* must carry a version of the endogenous essential gene that still retains function. Results of these crosses are summarized in Table 1 and presented in detail in Table S1-S3. As with *ClvR^tko^* (25), the combined female germline and maternal carryover-dependent cleavage to LOF for all four elements was very high (>99%); the male germline cleavage rate to LOF for the two new first generation *ClvR* elements was also high (from >94.7->99%). In short, the toxin component of the new *ClvR* elements is very efficient.

**Table 1.**
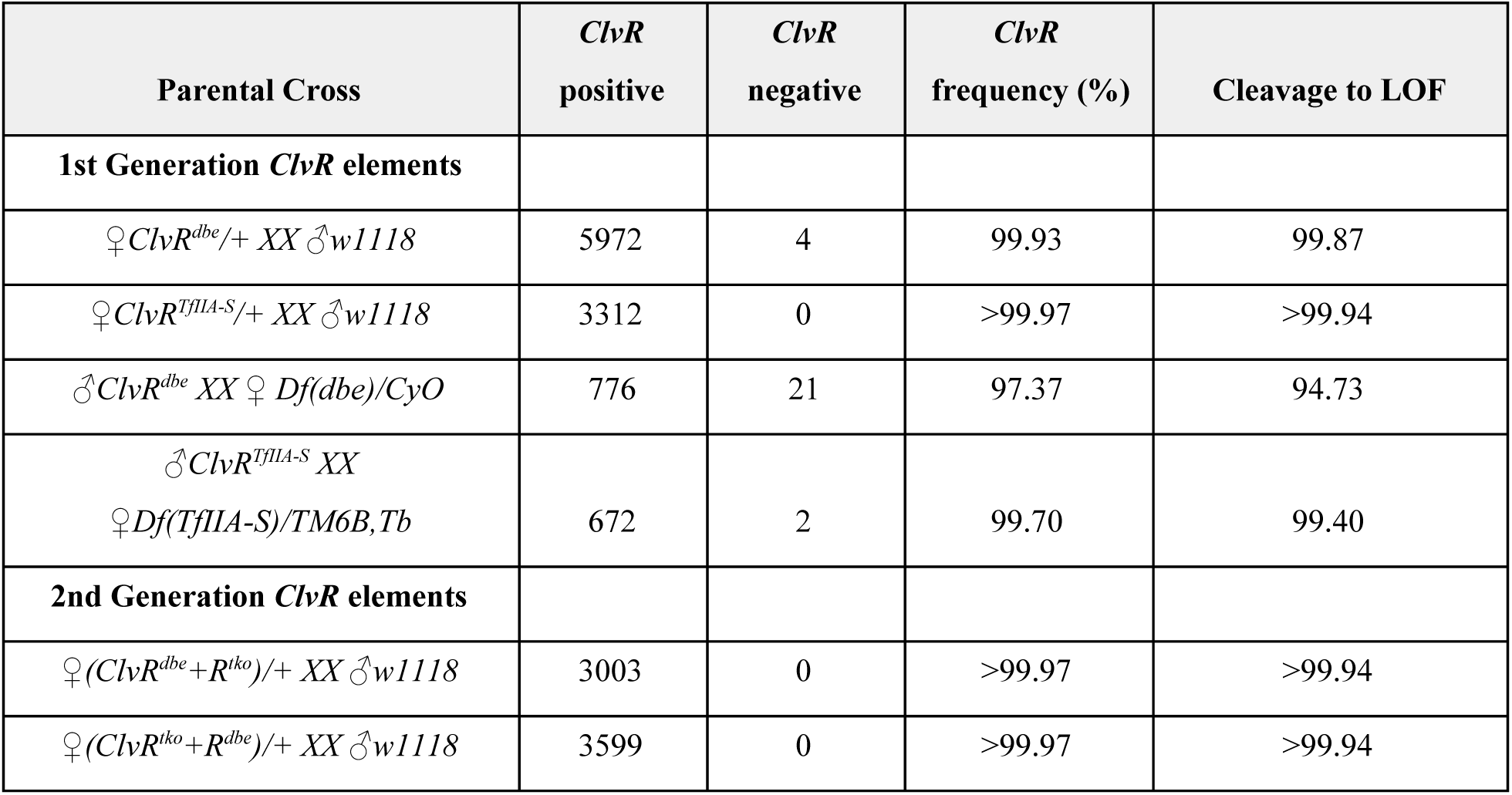
Genotype frequencies from crosses to determine the female and male cleavage rates to LOF.

### Analysis of target chromosomes following exposure to Cas9/gRNAs

For all flies from Table 1 that did not inherit a LOF allele we extracted genomic DNA and sequenced the target region. In addition, all male escapers were backcrossed to heterozygous *ClvR/+* females, and progeny scored for the absence of *ClvR*, to determine if the chromosome that escaped LOF allele creation was still sensitive to Cas9 cleavage and LOF allele creation. Details of the characterization are presented in Table S4 and Table S5. To summarize, all the escaper flies had at least 2 uncleaved wildtype target sites. Uncleavable target sites resulted mostly from small (3bp) in frame deletions or pre-existing polymorphisms and one rare 2bp substitution, all of which most likely preserved at least partial gene function. For all the escapers that we tested in a backcross to *ClvR/+* females, cleavage rates to LOF remained high (85 and 94% for two single cases, 100% for all the rest), indicating that the escaped chromosomes remained sensitive to *ClvR*-dependent LOF allele creation. Together these results are important because they provide further evidence that the *ClvR* approach to creation of LOF alleles utilizing four gRNAs is efficient and provides strong protection against the production of alleles at essential gene loci that have mutated target sites but retain essential gene function. Such resistant alleles can slow the rate of spread, and decrease the functional lifetime of *ClvR* elements in the population (25).

We also characterized cleaved target sites that were mutated to LOF following exposure to Cas9/gRNAs. In the case of *ClvR^tko^* all target sites could be cleaved, and a variety of indels were created (25). To explore these topics with *ClvR^dbe^* and *ClvR^TfIIA-S^* we characterized target sites following one generation of exposure to Cas9 and after 22 generations of a drive experiment (Fig. 3). The goal in looking at two different timepoints was to gain a sense of which target sites were preferentially cleaved, and if all target sites could be cleaved. In each case we carried out the analysis by taking *ClvR*-bearing flies and crossing them individually to flies that carried a deficiency for the target gene. From the progeny we selected one *ClvR*-bearing fly carrying a chromosome whose target locus had been exposed to Cas9/gRNAs, in trans to a deficiency to the region, and sequenced over the region between the four gRNA target sites. Sequencing results are summarized in Table S6-S9. For the *dbe* locus in *ClvR^dbe^* flies sites 3 and 4 were mutated at high frequency following a single generation of exposure to Cas9 (site 2 carried a SNP), and site 1 was altered in 3 of 16 analyzed flies. After 22 generations, however, all sites were altered. Products of cleavage included small deletions of varying sizes at single gRNA target sites, larger deletions between adjacent target sites, an inversion of the region between the outer target sites, and one fly had the whole locus deleted. In *ClvR^TfIIA-S^* the mutation spectrum was similar after 1 and 22 generations of exposure to Cas9. Most of the sequenced target sites had a deletion between gRNA 1 and 2 and smaller deletions at target sites 3 and 4 (19 and 26bp). Four flies had small deletions at each of the target sites and one fly had the whole region between gRNA 1 and 4 deleted. In summary, as with *ClvR^tko^*, all the target sites *ClvR^dbe^* and *ClvR^TfIIA-S^* could be cleaved, and a variety of different LOF indels were observed.

**Fig. 3:**
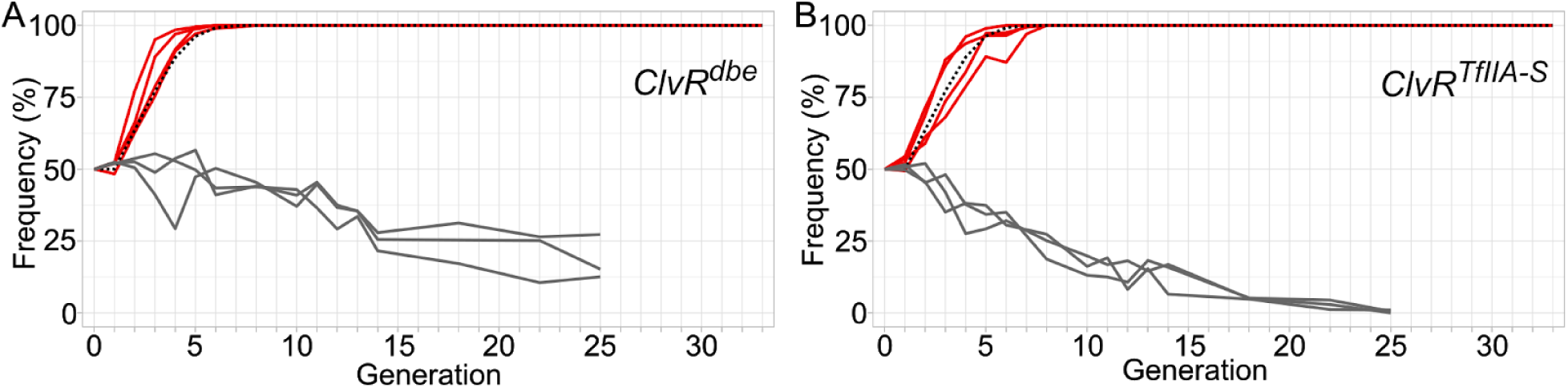
Gene drive experiments. Shown are the frequencies of *ClvR*-bearing flies (*ClvR*/+ and *ClvR*/*ClvR*) in a gene drive experiment with *dbe* **(A)** and *TfIIA-S* **(B)** as the targeted essential genes. Gene drive replicates are in red, modeled drive behavior for an element with no fitness cost in dotted black (see Methods for model details), and controls in grey.

### *ClvR^dbe^* and *ClvR^TfIIA-S^* spread to genotype fixation in wildtype *Drosophila*

To explore the behavior of the four new drive elements, *ClvR^dbe^*, *ClvR^TfIIA-S^, ClvR^tko^+R^dbe^, and ClvR^dbe^+R^tko^*, we carried out a set of six gene drive experiments, illustrated in Fig. 2, along with controls for each. We first tested the ability of *ClvR^dbe^* and *ClvR^TfIIA-S^* to spread in a wildtype population. As shown previously, *ClvR^tko^* spread to genotype fixation (>99.5% ClvR-bearing) within seven generations when introduced at a starting allele frequency of 25% (25) (and Fig. S6). The ability of *ClvR^dbe^* and *ClvR^TfIIA-S^* to spread was tested under similar conditions. The drive seed generation was set up by crossing male *ClvR*/+ heterozygous males to wildtype +/+ females, resulting in 50% of the population carrying one *ClvR* element in generation 1 (4 replicates, with a starting *ClvR* allele frequency of 25%). Control drive experiments, as with *ClvR^tko^*, were performed by crossing males heterozygous for a cassette that includes the *ClvR Rescue* and *td-tomato* marker, but not Cas9 and gRNAs, to wildtype +/+ females (3 replicates, with a starting control transgene allele frequency of 25%). Adult flies were allowed to lay eggs for one day in a food bottle. After approximately 14 days a large number of the eggs had developed into adults (∼700-1000). At that time point we sampled a random selection of the population and scored the frequency of the dominant *ClvR* (or control) marker. All the scored flies were transferred to a fresh food bottle to repeat the cycle. Results are plotted in Fig. 3. Both *ClvR^dbe^* and *ClvR^TfIIA-S^* reached genotype fixation (100% of flies having 1 or 2 copies of *ClvR^dbe^*) between 6 and 8 generations in all replicates. The control elements slowly decreased in frequency over time, perhaps due to some fitness cost associated with the ubiquitous expression of a dominant marker gene and/or the presence of additional copies (for a total of 3-4 in transgene-bearing individuals) of the essential target gene present in the *Rescue* only control element.

After 22 generations (32 generations for *ClvR^tko^*) we assayed the allele frequencies in the drive populations (Table S10). These ranged from 83% to 90% for *ClvR^dbe^*, 91% to 96% for *ClvR^TfIIA-S^*, and 94 to 100% for *ClvR^tko^* populations. These observations demonstrate that multiple *ClvR* elements, each targeting a different gene, can be generated and show gene drive when located at a common site in the genome.

### 2nd generation *ClvR-*elements *ClvR^tko^+R^dbe^* and *ClvR^dbe^+R^tko^* spread to genotype fixation

Second generation *ClvR* elements face several potential challenges to spread in wildtype and *ClvR^n^*-bearing populations. First, they carry an additional cargo in the form of the *ClvR^n^* Rescue transgene, which may introduce a fitness cost, particularly when drive occurs into a wildtype background in which the *ClvR^n^ Rescue* does not function to support drive (it results in *ClvR*-bearing individuals having 3 or 4 functional copies of essential gene^n^). Second, when driving into a population of *ClvR^n^*, both *Rescue* transgenes must work well and be able to rescue LOF phenotypes for two essential genes. Finally, when driving into a population of *ClvR^n^*, *ClvR^n+1^* elements will also often find themselves in a transheterozygous state with the earlier generation element. These *ClvR^n^* elements include a different set of four gRNAs that do not contribute to drive by *ClvR^n+1^*. These could compete with the four gRNAs needed for drive by *ClvR^n+1^* for loading into a complex with Cas9, thereby suppressing drive. To explore the ability of *ClvR^n+1^* elements to thrive in different genetic backgrounds we carried out drive experiments of *ClvR^n+1^* elements into wildtype and *ClvR^n^*-bearing populations. Drive experiments of *ClvR^tko^+R^dbe^* and *ClvR^dbe^+R^tko^* into wildtype *w^1118^* populations were carried out as with *ClvR^dbe^* and *ClvR^TfIIA-S^* above, by mating heterozygous *ClvR*/+ males to *w^1118^*; +/+ virgins, for a starting *ClvR* allele frequency of 25%. As a control for these experiments we also carried out drive into populations fixed for a *ClvR* element carrying the same *Rescue* transgene as that needed for drive by *ClvR^n+1^* (drive of *ClvR^tko^+R^dbe^* into *ClvR^tko^*, and drive of *ClvR^dbe^+R^tko^* into *ClvR^dbe^*). For the experiments involving drive into populations of *ClvR^dbe^* and *ClvR^tko^* (the controls above, and drive of *ClvR^tko^+R^dbe^* and *ClvR^dbe^+R^tko^* into *ClvR^dbe^* and *ClvR^tko^*, respectively) we crossed heterozygous *ClvR^n+1^*/+ males to virgin females taken from the *ClvR^dbe^* and *ClvR^tko^* drive populations at the generation where we determined allele frequency in the drive populations (generation 22 for *ClvR^dbe^*, Fig. 3, and Generation 32 for *ClvR^tko^* Fig. S6). These populations consist of mostly *ClvR*/*ClvR* homozygotes, with a few *ClvR*/+ individuals (Table S 10).

The outcomes of these gene drive experiments are shown in Fig. 4. *ClvR^dbe^+R^tko^* reached genotype fixation between 7 and 8 generations when driving into a population of wildtype *w^1118^*; +/+, and between 8 and 9 generations when driving into a population of *ClvR^tko^*. As expected, when driving into a population of *ClvR^dbe^*, the 2nd generation *ClvR^dbe^+R^tko^* drive element did not increase in frequency (Fig. 4A). *ClvR^tko^+R^dbe^* performed similarly, though with slightly slower kinetics. *ClvR^tko^+R^dbe^* reached genotype fixation between 8 and 10 generations when driving into a population of *w^1118^*, and between 9 and 10 generations when driving into populations of *ClvR^dbe^*. When driving into a population of *ClvR^tko^*, *ClvR^tko^+R^dbe^* did not increase in frequency (Fig 4B). For the experiments in which a *ClvR^n+1^* was driven into a population of *ClvR^n^* (*ClvR^dbe^+R^tko^* into *ClvR^tko^*, and *ClvR^tko^+R^dbe^* into *ClvR^dbe^*) we also measured the allele frequency of the *ClvR^n^* elements at generation 12. As expected based on the fact that *ClvR^n^* and *ClvR^n+1^* share a common genomic location, the high frequency of *ClvR^n+1^* at genotype fixation, with allele frequencies ranging from 84.9-91.2%, was associated with a dramatic decrease in the allele frequency of *ClvR^n^*, to between 8.2-14.5% (Details in Tables S11 and S12). Together these results demonstrate that second generation *ClvR* elements can be created; they drive themselves into wildtype populations; they also drive into populations fixed for an earlier generation element, and they displace the latter as they spread. Thus, cycles of gene drive-mediated population modification can be achieved, while at the same time bringing about a decrease in the frequency of an earlier generation element.

**Fig. 4:**
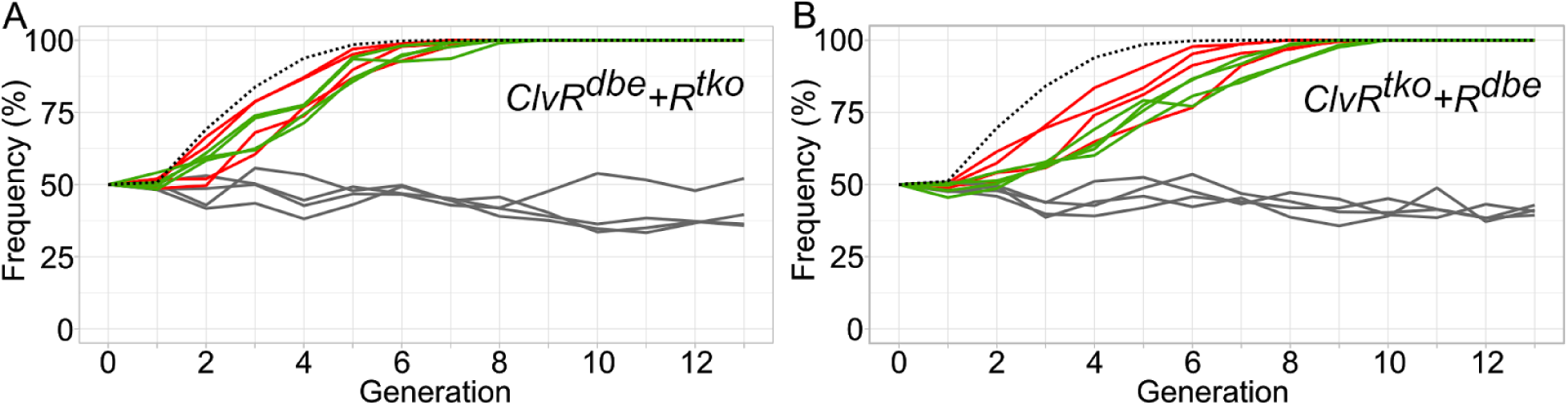
*ClvR^n+1^+R^n^* selfish elements drive into populations carrying *ClvR^n^*. Shown are the drive outcomes for **(A)** *ClvR^dbe^+R^tko^* driving into a population of *w^1118^* (red lines), *ClvR^tko^* (green lines), and *ClvR^dbe^* (grey lines, control), and **(B)** *ClvR^tko^+R^dbe^* driving into a population of *w^1118^* (red lines), *ClvR^dbe^* (green lines), and *ClvR^tko^* (grey lines, control) There are four replicates for each drive experiment. Dashed black lines represent model estimated behavior for the spread of *ClvR*^n+1^ with no fitness cost into populations consisting primarily of *ClvR*^n^ population (see Methods for model details). Allele frequencies at generation 12 are presented in Table S11-12

## Discussion

Genetic modification of a population is unlikely to ever be a one shot project, with a single genetic element providing all desired modifications, for an indefinitely long time. Mutation, recombination and natural selection will cause a loss of drive and/or efficacy through one mechanism or another. In addition, as knowledge increases there are likely to be situations in which one wants to augment (upgrade) a population modification, or remove a change whose presence is no longer desired. For all these reasons it is important that strategies for population modification be extensible for multiple cycles. At the same time, it is also important that introductions of new elements result in the loss (or at least a great decrease in the frequency) of old elements from the population.

Here we show that a first set of cycles of modification with these properties can be achieved using *ClvR* selfish genetic elements. While modeling is required to provide a detailed analysis, we can intuit several ways in which it may be possible to carry out cycles of modification indefinitely: as a linear chain of elements, in which the newest element always carries the *Rescue* of the element from the previous generation, with each new element cleaving and rescuing a new essential gene, or as a circle in which variants of elements that target in total three or more different genes are used repeatedly.

In order to achieve multiple cycles of population modification the drive components need to be orthogonally acting and indefinitely extensible. The components of *ClvR* are orthogonally acting since Cas9/gRNAs and *Rescue* transgenes are specific to a particular essential gene. *ClvR* is also in principal broadly extensible within a species since any gene that is essential for survival or fertility can be targeted for cleavage and rescue. In our earlier work, we created *ClvR^tko^*, which is located on the third chromosome and targets an essential gene on the X. Here we showed that additional *ClvR*s can be generated at the same site as *ClvR^tko^*, targeting essential genes involved in different biological processes, located on chromosomes 2 and 3. Further evidence for the extensibility of the *ClvR* system comes from recent work in which it was shown that a construct consisting of two gRNAs targeting the gene encoding the essential developmental transcription factor *hairy* (*h*) and a recoded rescuing version of *h* could, when located within the *h* locus, spread through a population homozygous for Cas9 at an independent locus through a *ClvR*-like mechanism (22). Spread to transgene fixation occurred rapidly for all four of these elements. Together these observations show that *Cleave and Rescue* type selfish genetic elements can successfully mutate to LOF and provide rescuing essential gene function for genes involved in a variety of cellular processes, and that no particular spatial relationship in the genome between the drive element and target gene is essential. We also note that while our work herein and in (22, 25) used Cas9, which generates LOF mutations by creating double strand DNA breaks that are repaired inaccurately, similar LOF effects can be brought through any mechanism that modifies DNA site-specifically, including methods that do not create double-strand breaks. Examples of other possible mechanisms include a Cas9/gRNA-linked base editor (33), a Cas9/gRNA nickase linked to reverse transcriptase (34), or a pair of site-specific engineered recombinases (35).

For the two *ClvR*s we generated here, and the one in (25), a variety of indels were created in the essential genes targeted, and all sites were ultimately cleaved. Importantly, in our work and that of Champer and colleagues (22) alleles of the endogenous essential gene that were completely resistant to cleavage but retained function were not observed. We note that in (25) and the current work examples were found in which two of the four gRNA target sites were altered in ways that presumably retained function. In some cases one of these was due to a preexisting polymorphism that was not screened for prior to initiating these experiments. However, other sequence differences were new and due to mutation associated with inaccurate DNA repair. Thus, we recommend multiplexing no less than four gRNAs to maintain functionality in genetically diverse populations. The feasibility of using more than four gRNA to bring about increased robustness in terms of LOF allele creation is suggested by our observation that *ClvR^n+1^* elements showed drive even when in the presence of the four additional gRNAs present in *ClvR^n^*, which do not contribute to drive (Fig. 4).

*ClvR*-type gene drives can modify populations in a number of ways. In one family of approaches *ClvR* spreads a LOF allele through the population. This can happen if *ClvR* is itself located within a gene of interest, thereby disrupting it and driving an increase in frequency of the disrupted allele as it spreads. Alternatively the *ClvR* can carry gRNAs that target some other gene for LOF allele creation following cleavage. In a second family of approaches *ClvR* can carry into a population linked cargo transgenes, or an allele of an endogenous locus to which it is tightly linked through insertion site choice. In all of these scenarios drive only occurs when the fitness costs to non-*ClvR*-bearing individuals exceed those associated with being *ClvR*-bearing, attributes that are frequency-dependent (25, 30). Finally, while more speculative, we note that the mechanism by which *ClvR*-based population modification occurs provides some unique opportunities for strategies that prevent disease transmission by bringing about the death of host cells and/or hosts in response to infection, or that bring about periodic overall population suppression in response to a cue from the environment. These ideas each take advantage of a key feature of *ClvR*-dependent drive – that individuals of the modified population are absolutely dependent on the functionality of the *Rescue* transgene for survival. Given this it is interesting to imagine ways in which the function of the *Rescue* transgene could be made conditional so as to bring about the death of cells, individuals or populations under specific circumstances. For example, it may be possible to engineer essential gene function at the level of transcript or protein such that it is sensitive to the presence of viral protease activity (4), small RNAs (36), or other honest markers of infection, resulting in the death of infected host cells or individuals. One can also imagine ways in which entire *ClvR*-bearing populations could be suppressed in an environmental condition-specific manner. Temperature, to give an example, is often an important seasonal environmental variable. Gene function can be made temperature sensitive in several ways. A temperature sensitive intein can be incorporated into the coding region of the essential gene (37). Because self-splicing from the encoded protein is temperature dependent, survival of a population in which all wildtype alleles are LOF should be so as well. Alternatively, a temperature sensitive degron could be linked to the coding region for a similar effect (38). Strategies for engineering essential gene function to be sensitive to the presence of specific chemicals could use a similar strategy, in which protein degradation occurs in response to binding of a small molecule ligand to a specific protein domain (39–41).

One can imagine scenarios in which each of the above approaches to population modification with *ClvR* is successful and efficacious. However, a similar analysis of each will also identify multiple mechanisms by which drive and efficacy of any cargo can fail over time. This does not mean that population modification should not be attempted. Interventions in any area of biology that involves the forces of mutation and selection typically come with periods of success followed by failure. The evolved resistance of cancers to specific therapies, of bacteria to antibiotics and phage therapy, of plasmodium to antimalarial drugs, of insects and plants to insecticides and herbicides, and the yearly battle of the human immune system and the latest vaccines against the current strain of influenza all reflect the ubiquitous nature of this cycle. The important thing is to have a plan that allows for the continual evolution and implementation of an initially successful strategy. In the case discussed herein, where the goal is to alter the genetic composition of a population toward a specific functional end, the ability to iteratively carry out new modifications while removing old ones provides the essential underpinnings of any plan for long-term success.

## Methods

Restriction Enzymes, Gibson Assembly enzymes, Q5 and Longamp DNA polymerases were from NEB. Genomic DNA extraction kit (*Quick*-DNA 96 Plus Kit), mini plasmid prep (ZymoPURE Plasmid Miniprep), and gel extraction kit (Zymoclean Gel DNA Recovery Kit) were from Zymo Research. DNA maxiprep kit was from Qiagen (EndoFree Plasmid Maxi Kit). All plasmids were cloned with Gibson assembly (42) as described previously (25). All fly embryonic injections were performed by Rainbow Transgenic Flies. Cloning construct design, CRISPR guide design (43) and sequencing alignments with MAFFT (44) were done in the Benchling software suite. All primers, gRNA target sequences, and construct fasta files are in Dataset S1.

### Cloning of *ClvR* constructs and generation of 1st generation *ClvR* flies

1st generation *ClvR* flies (*ClvR^dbe^* and *ClvR^TfIIA-S^*) were generated in two steps as described previously (25). We first inserted the *Rescue* part into the fly genome, followed by integration of the *Cleaver* (Cas9 and gRNAs) at that same site. The first construct had the *Rescue* of the target gene which was amplified from genomic DNA of *D. suzukii* (*Dsuz*). The *Rescue* fragments contained the ORF of the target gene as well as upstream and downstream sequences with potential promoter/enhancer and terminator elements. The *Dsuz-dbe Rescue* fragment was 2kb, the *Dsuz-TfIIA-S* fragment was 3.8 kb (annotated fasta files in Fig S7-8). In addition, the construct had an *opie-td-tomato* dominant marker and an *attP* site. All these elements were flanked by homology arms to facilitate CRISPR mediated homologous recombination (HR) into the fly genome. Outside the homology arms the constructs had a U6 driven gRNA that targeted a site at 68E on the 3rd chromosome of *Dmel* (Fig S1A).

The constructs were injected into a stock that had *nos-Cas9* on the X-chromosome (Fig. S2A) (45). G0 injected flies were outcrossed to *w^1118^* and the progeny screened for ubiquitous *td-tomato* expression. Male transformants that came from a male G0 fly were outcrossed again to *w^1118^* to build up a stock. At this point the Cas9 source on the X-chromosome of the injection strain was bred out.

The second part of the *ClvR* element (the *Cleaver*) was assembled separately and had Cas9 driven by germline specific *nos* promoter and UTRs (46) (*nos*-Cas9 derived from addgene plasmid #62208 (45)). A set of 4 gRNAs were each driven from alternating pairs of U6:3 and U6:1 promoters (28) (similar as in (45). The plasmid further had an *attB* site to facilitate integration into the genomic location of the first construct, and a *3xP3-*GFP transformation marker (Fig. S1B).

This construct was injected into flies that carried the *Rescue* alongside a helper plasmid as a source of *phiC31* integrase. G0 injected flies were outcrossed to *w^1118^* and screened for eye-specific expression of GFP (See Fig. S2B).

### Generation of 2nd Generation *ClvR^n^+R^n-1^* flies

We used CRISPR mediated homologous recombination to modify the *ClvR* locus of the strain of flies that would become *ClvR^n+1^*. A Cas9/gRNA ribonucleoprotein (RNP) complex binding between the *3xP3* promoter and the GFP ORF at the original *ClvR* locus was injected into *ClvR* flies alongside a donor plasmid to be inserted via CRISPR HR (Fig. S2C). The RNP complexes were assembled by mixing Cas9 protein (Alt-R, IDT) and gRNA (sgRNA, IDT) in water and incubating for 5 min at room temperature. Afterwards the donor plasmid was added and the mixture was stored at −80℃ until injection. Final concentrations in the injection mix were: Cas9 protein 500ng/ul, gRNA 100ng/ul, donor plasmid 500ng/ul. The donor plasmid contained the *Rescue* of *ClvR*^n^ and an *opie2* promoter (47) with partial GFP sequences that acted as the homology arm. The other homology arm was the *3xP3* promoter and plasmid backbone (Fig S1C). We injected a construct carrying the *D. suzukii-dbe Rescue* into *ClvR^tko^* flies to generate 2nd generation *ClvR^tko^+R^dbe^* flies. We also injected a construct carrying the *Drosophila virilis* derived *tko Rescue* into *ClvR^dbe^* flies to create *ClvR^dbe^+R^tko^* flies. Successful integration of the *Rescue*^n^ construct was detected by ubiquitous GFP expression. To confirm the integration of the new *Rescue* at the correct genomic location we extracted genomic DNA from GFP positive flies and amplified a fragment with primers binding in the *3xP3* promoter and the nanos 3’UTR downstream of Cas9. The resulting PCR fragments were partially sequenced to confirm that the new *Rescue* was downstream of *3xP3* and that *opie*-GFP was downstream of the nos 3’UTR (Fig. S2C).

### Crosses to determine male and female cleavage rates to LOF

We crossed *ClvR*-bearing males to *w^1118^* virgins to get heterozygous *ClvR/+* male and female offspring. To determine the female cleavage rate to LOF we took *ClvR/+* virgins, crossed them to *w^1118^* males, and scored the progeny for the *ClvR* marker *td-tomato*. *ClvR* frequency was calculated as number of *td-tomato* flies divided by the total number of flies (Table S1). The cleavage rate to LOF is *ClvR*-positive progeny divided by half the total progeny, since with mendelian inheritance 50% of the progeny would be expected to inherit *ClvR* in the absence of *ClvR* dependent killing. The same cross was performed with 2nd Generation *ClvR^n+1^+R^n^* flies (Table S3). For the male *ClvR* frequency we crossed heterozygous *ClvR*/+ males to a stock that carried a deficiency for the target gene. *ClvR* frequency was calculated by determining the fraction of the total carrying the *Df (target essential gene)* that were also *ClvR*-bearing (Table S2).

### Sequencing analysis of escapers and cleavage events

Whenever possible we isolated the chromosome that we wanted to sequence over a *Df* for the essential gene so that there was only one version of the essential gene available. This was done for all sequenced flies except for the 4 escapers coming from heterozygous *ClvR^dbe^/+* females (Table S4-5). Genomic DNA was extracted with the *Quick*-DNA 96 Plus Kit from Zymo. For *dbe* we amplified a 2.2kb genomic region spanning all target sites with primers dbe-genomic-F and dbe-genomic-R and Sanger-sequenced that amplicon with primers dbe-seq-F and dbe-seq-R. For *TfIIA-S* we amplified a 1.9kb genomic region with primers tf2-genomic-F and tf2-genomic-R. This amplicon was Sanger-sequenced with primer tf2-genomic-R. See Fig. S3 for a schematic of the genomic regions and primer binding sites.

### Gene drive experiments

All *ClvR* drive experiments were set up as described previously (25). We crossed heterozygous *ClvR/+* males to *w^1118^* virgins in bottles of fly food. After 2 days the adults were removed and the progeny (seed generation=0, *ClvR*-bearing=50%, allele frequency=25%) were allowed to eclose in the bottles. After 13-14 days, eclosed flies were anesthetized on a CO2 pad and a random sample of approximately 200 flies was scored for the dominant *ClvR* marker. This sample was then transferred to a bottle with fresh food to continue the next generation. All counts of gene drive experiments are in Dataset S2.

### *ClvR* Computational Model

Figures 3 and 4 feature model predicted behavior for a *ClvR* driving into wildtype, or a 2nd generation *ClvR^n+1^* driving into a population fixed for a 1st generation, *ClvR^n^*, with 0% fitness costs as well as 100% cleavage and maternal carryover rates. We used a deterministic, population proportion model adjusted from a model we have used previously (25) which uses difference equations to track the frequency of each genotype over discrete generations. In this model we assumed that there is random mating; females produce offspring from a single mating; cleavage occurs during gametogenesis; maternal carryover of Cas9 and gRNAs can cleave any uncleaved allele in the zygote, such as that coming from the father; being heterozygous for a cleaved allele has no fitness effects (the locus is haplosufficient), and two copies of the cleaved target without a *Rescue* results in death 100% of the time.

### Fly crosses and husbandry of *ClvR*^tko^ flies

Fly husbandry and crosses were performed under standard conditions at 26°C. Rainbow Transgenic Flies (Camarillo, CA) carried out all of the embryonic injections for germline transformation. Containment and handling procedures for *ClvR* flies were as described previously (28), with G.O and B.A.H. performing all fly handling.

## Data availability

All data is available in the main text and the supplementary materials. *ClvR* flies are available on request under the conditions outlined in (25).

## Acknowledgments

Stocks obtained from the Bloomington Drosophila Stock Center (NIH P40OD018537) were used in this study.

## Contributions

Conceptualization, G.O., T.I. and B.A.H.; Methodology, G.O., T.I. and B.A.H.; Investigation, G.O. and B.A.H.; Modeling, T.I.; Writing – Original Draft, G.O. and B.A.H.; Writing – Review & Editing, G.O., T.I. and B.A.H.; Funding Acquisition, G.O. and B.A.H.

## Supplementary Information for

Other Supplementary Materials for this manuscript include the following:

Dataset S1. DataS1.xlsx; primer sequences, gRNA sequences, ClvR constructs fasta files

Dataset S2. DataS2.xlsx; genotype count table for drive experiments.

### Supplementary Figures

**Fig. S1.**
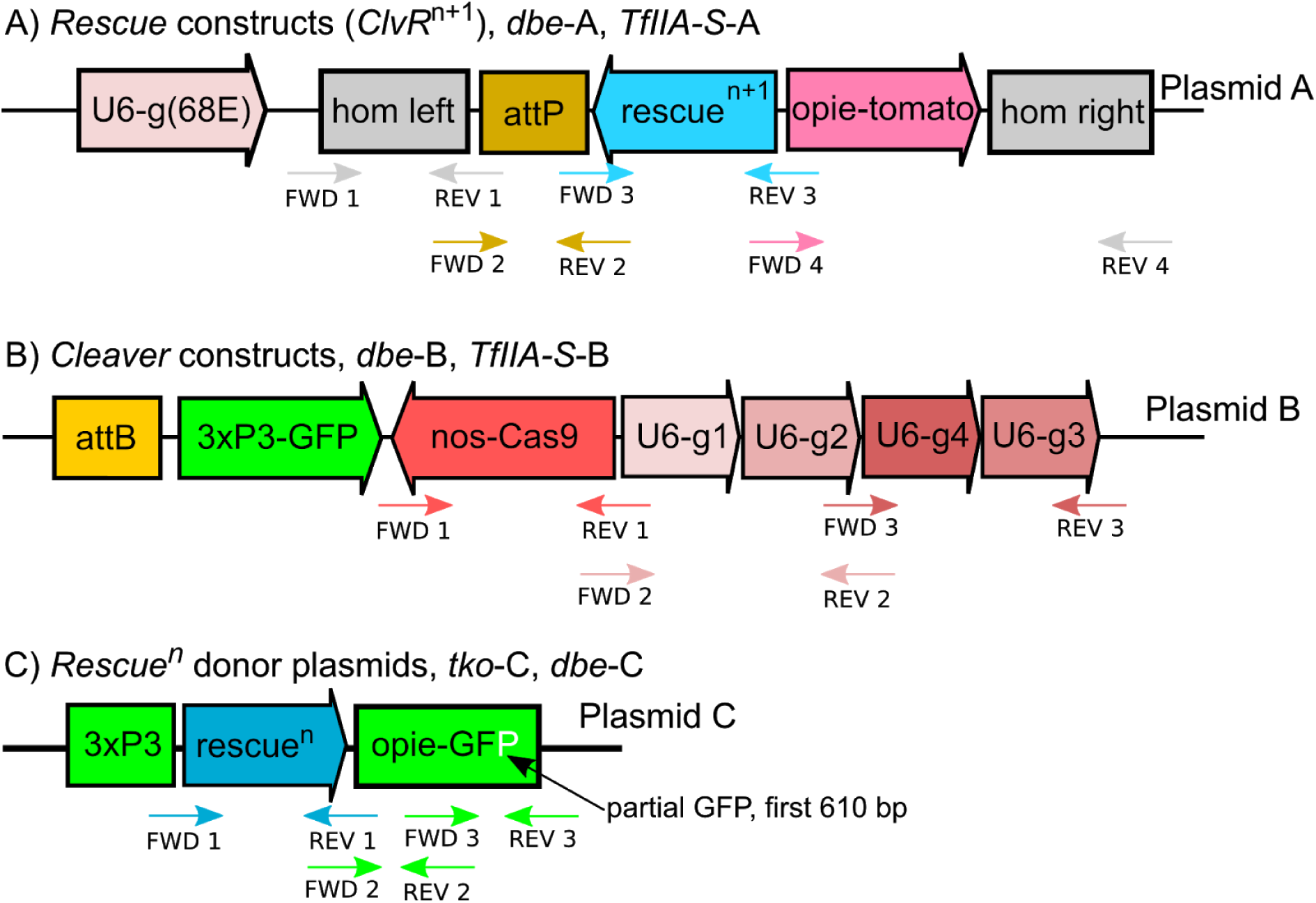
Constructs used to create 1st and 2nd generation *ClvR* flies. Schematic representation of *ClvR* constructs including primers used for cloning. Primer sequences are in Dataset S1. **(A)** *Rescue* constructs including the *Rescue* gene, a *td-tomato* marker and an *attP* site, flanked by homology arms to facilitate insertion into the fly genome at 68E on the third chromosome. The gRNA to target 68E was driven from a U6 promoter located outside of the homology arms. **(B)** *Cleaver* constructs having an *attB* site, a *3xP3-*GFP marker gene, germline Cas9 under the control of the *nanos* promoter and UTRs, as well as a set of 4 gRNAs each driven by a U6 promoter. **(C)** *Rescue^n^* donor plasmid having a *3xP3* promoter serving as homology arm, the *Rescue^n^* gene, a ubiquitous *opie* promoter and a partial GFP ORF (110 bp +UTR (SV40) missing the C-terminus of GFP served as the other homology arm.

**Fig. S2.**
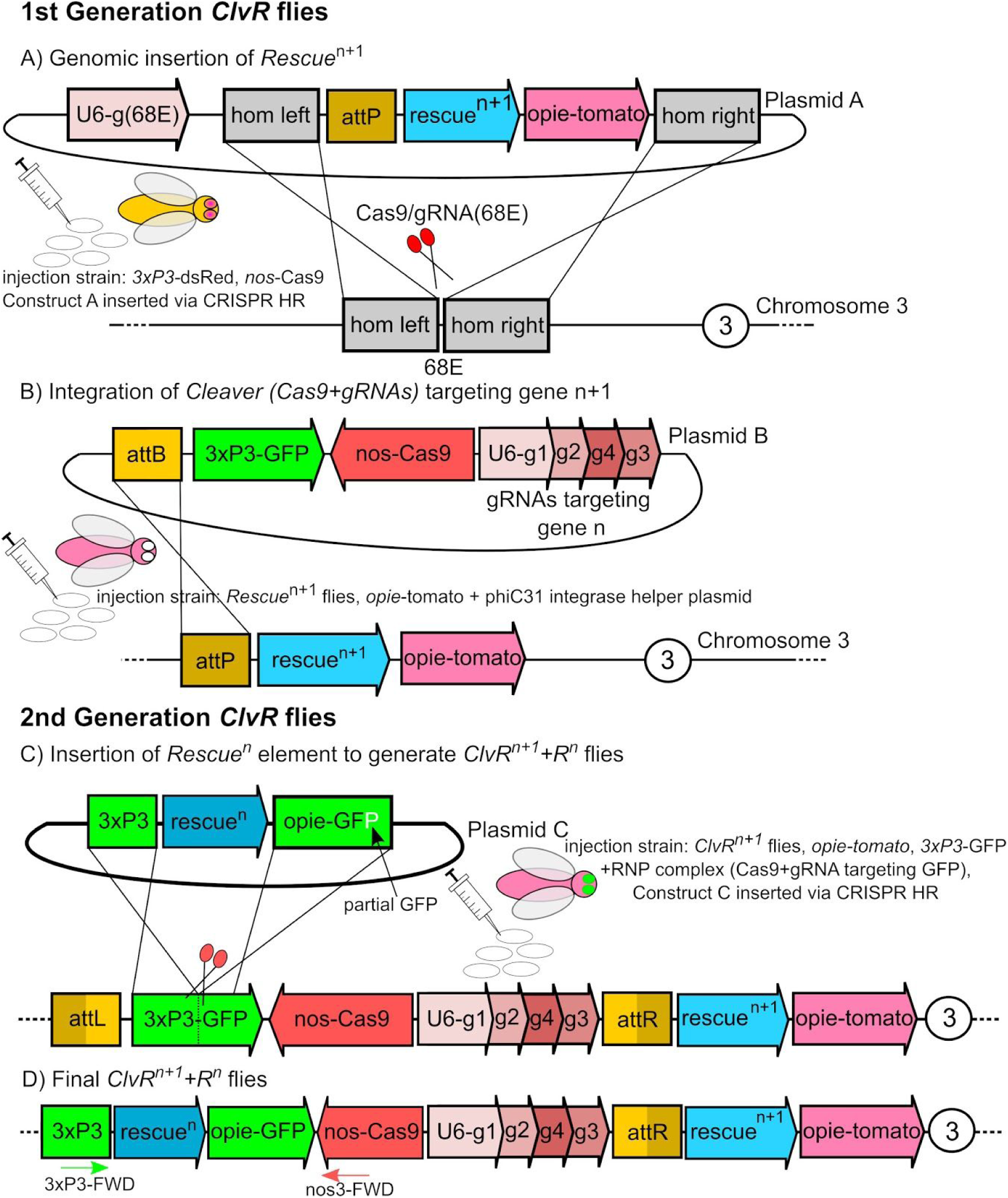
Synthesis strategy to create 1st and 2nd generation *ClvR* flies. **(A) CRISPR HR mediated insertion of *Rescue^n+1^*.** Plasmid A having the *Rescue^n+1^* and a marker was injected into a strain expressing Cas9 in the germline (*nos*-Cas9). A gRNA targeting a genomic region at 68E was expressed from the plasmid outside the homology arms. **(B) *PhiC31* mediated integration of *Cleaver^n+1^* (Cas9 and gRNAs).** Plasmid B having Cas9 and gRNAs targeting essential gene (n+1) was injected into flies from step A with a helper plasmid as the source for *phiC31* integrase. **(C) CRISPR HR mediated insertion of *Rescue^n^* into flies that will become *ClvR^n+1^+R^n^* flies.** Cas9/gRNA RNP complexes were injected to induce a DSB between the *3xP3* promoter and GFP alongside a donor template that had the *Rescue^n^*. The homology arms were designed in a way so that successful insertion will switch GFP expression from eye-specific to ubiquitous. **(D) Final *ClvR^n+1^+R^n^* flies.** These flies were used in the gene drive experiments to replace populations carrying *ClvR^n^* elements. Red and green arrows indicate primers that were used to confirm correct insertion of the new *Rescue^n^*.

**Fig. S3.**
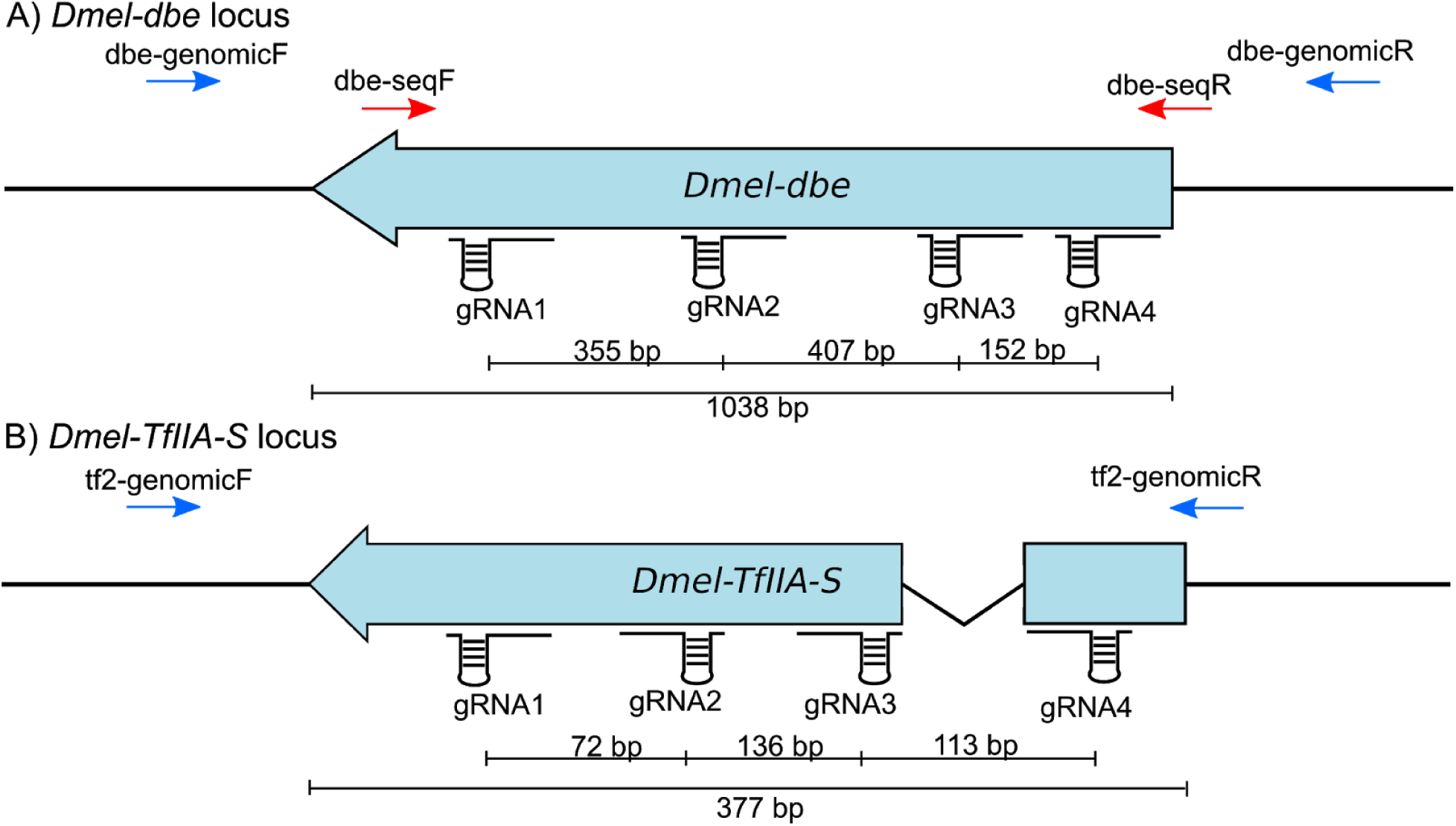
Schematic of target genes genomic loci. Shown are exons (blue) and introns of the two *ClvR* target genes with gRNA binding sites and directions, primers used for sequencing, and scale bars giving distances in bp. **(A)** *Dmel-dbe* locus **(B)** *Dmel-TfIIA-S* locus

**Fig. S4.**
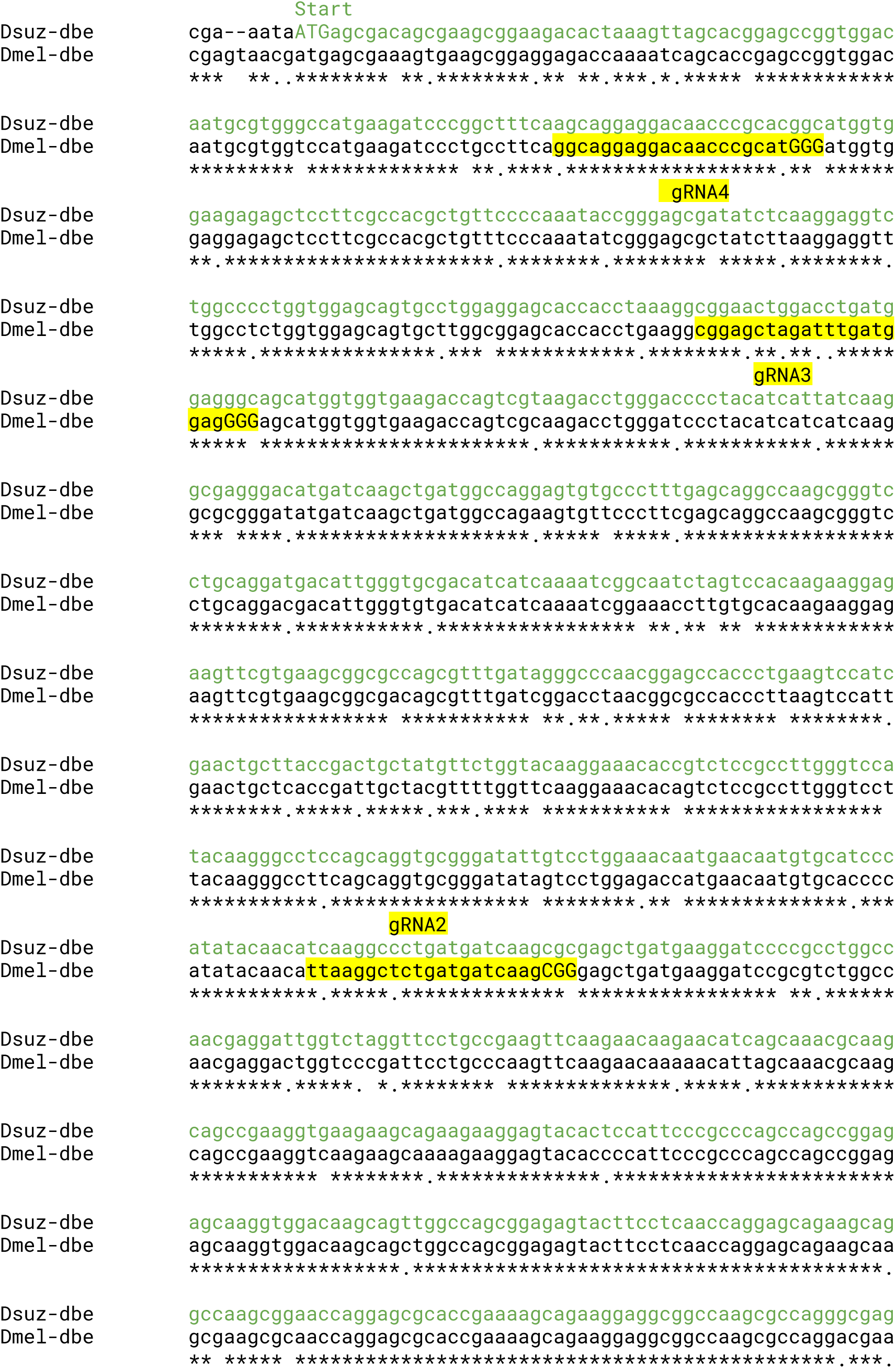
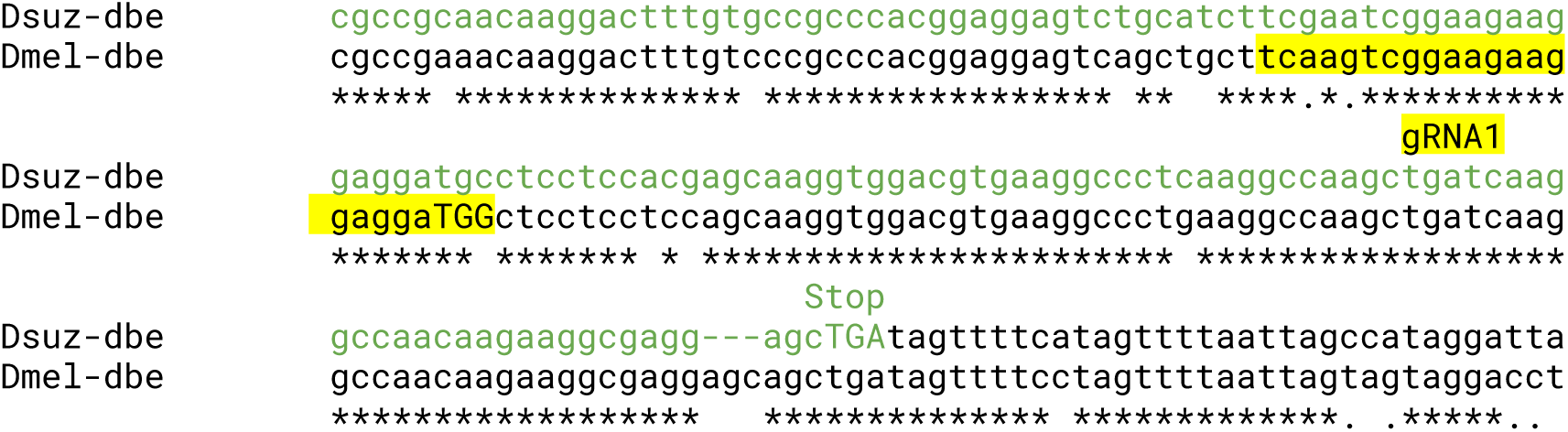
Alignment of *dbe* target region with the *Rescue* fragment. Shown is the DNA sequence alignment of the *dbe* locus gRNA target sites in *D. melanogaster* with the *Rescue* fragment form *D. suzukii*. Note how the gRNAs can only target the *D. melanogaster* locus. CDS in green, Start/Stop in uppercase, gRNA target sites including PAM in yellow, PAM in uppercase.

**Fig. S5.**
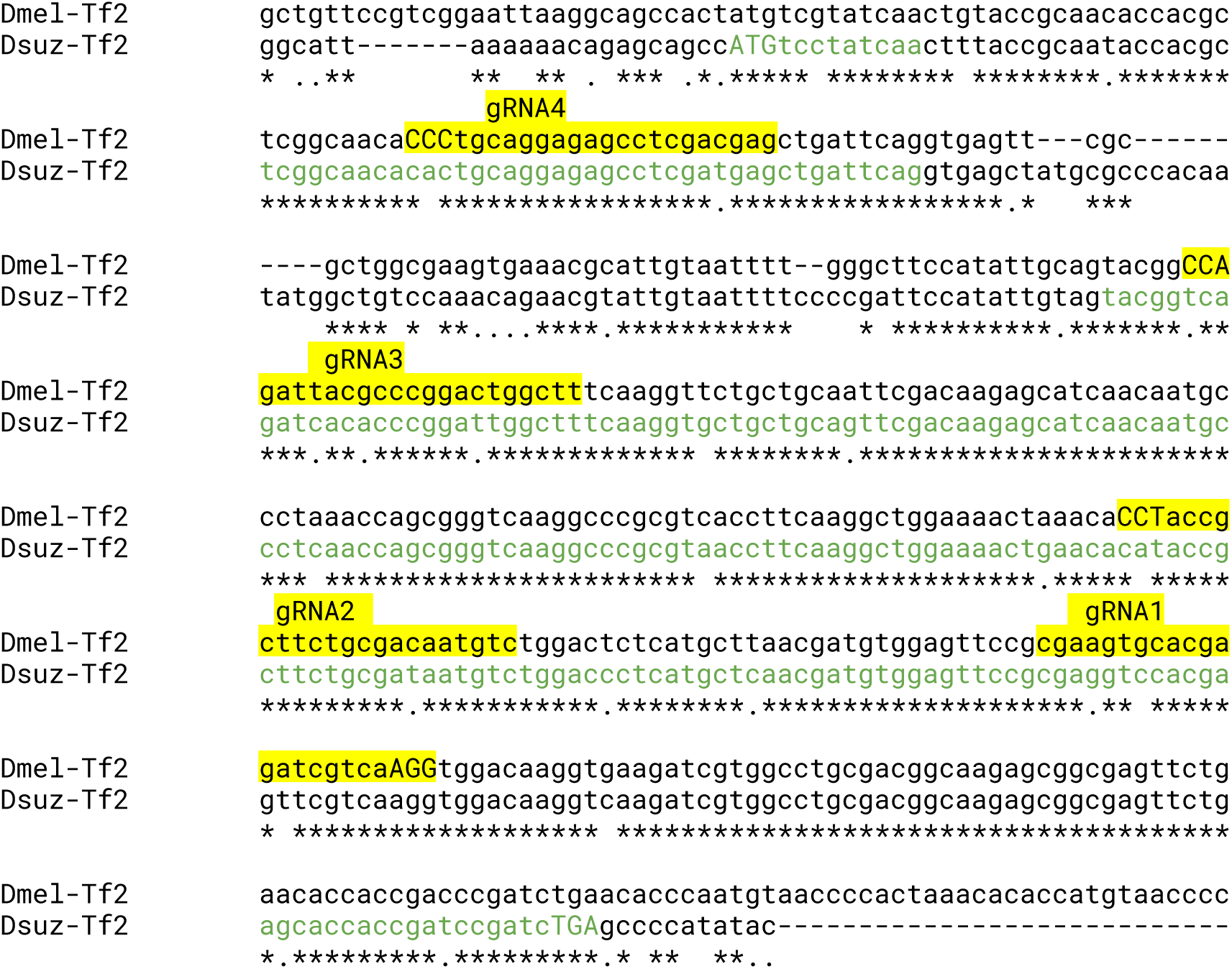
Alignment of *TfIIA-S* target region with the *Rescue* fragment. Shown is the DNA sequence alignment of the *TfIIA-S* locus gRNA target sites in *D. melanogaster* with the *Rescue* fragment from *D. suzukii*. Note how the gRNAs can only target the *D. melanogaster* locus. CDS in green, Start/Stop in uppercase, gRNA target sites including PAM in yellow, PAM in uppercase.

**Fig. S6.**
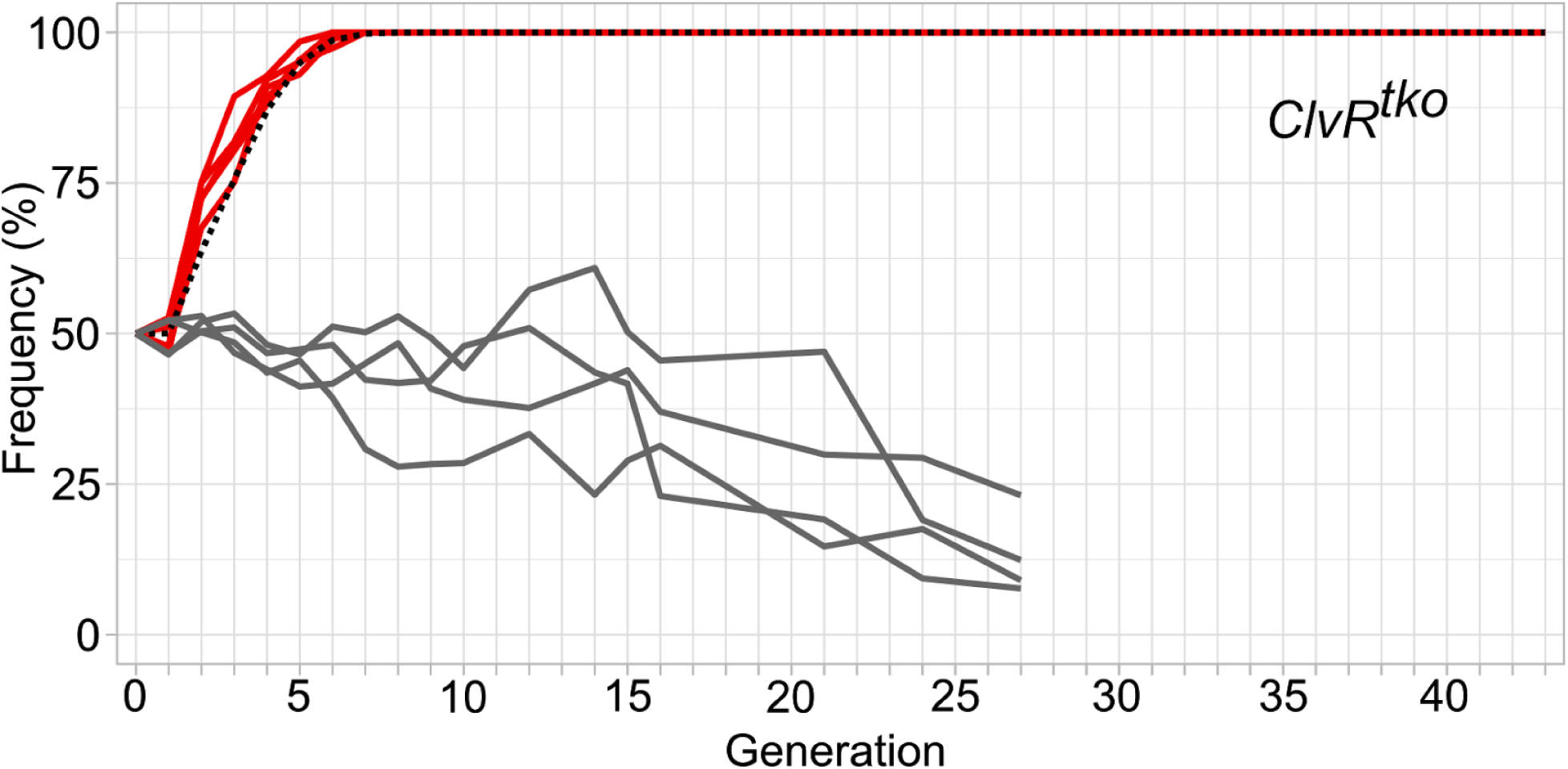
Continuation of a *ClvR^tko^* gene drive experiment reported in (25). Red lines represent drive data from five replicates. Grey lines show data from control experiments in which a construct carrying only the *tko Rescue* and a dominant marker was introduced into a *w^1118^* population. Dotted black line shows model behavior for an element with no fitness cost.

**Fig S7.**
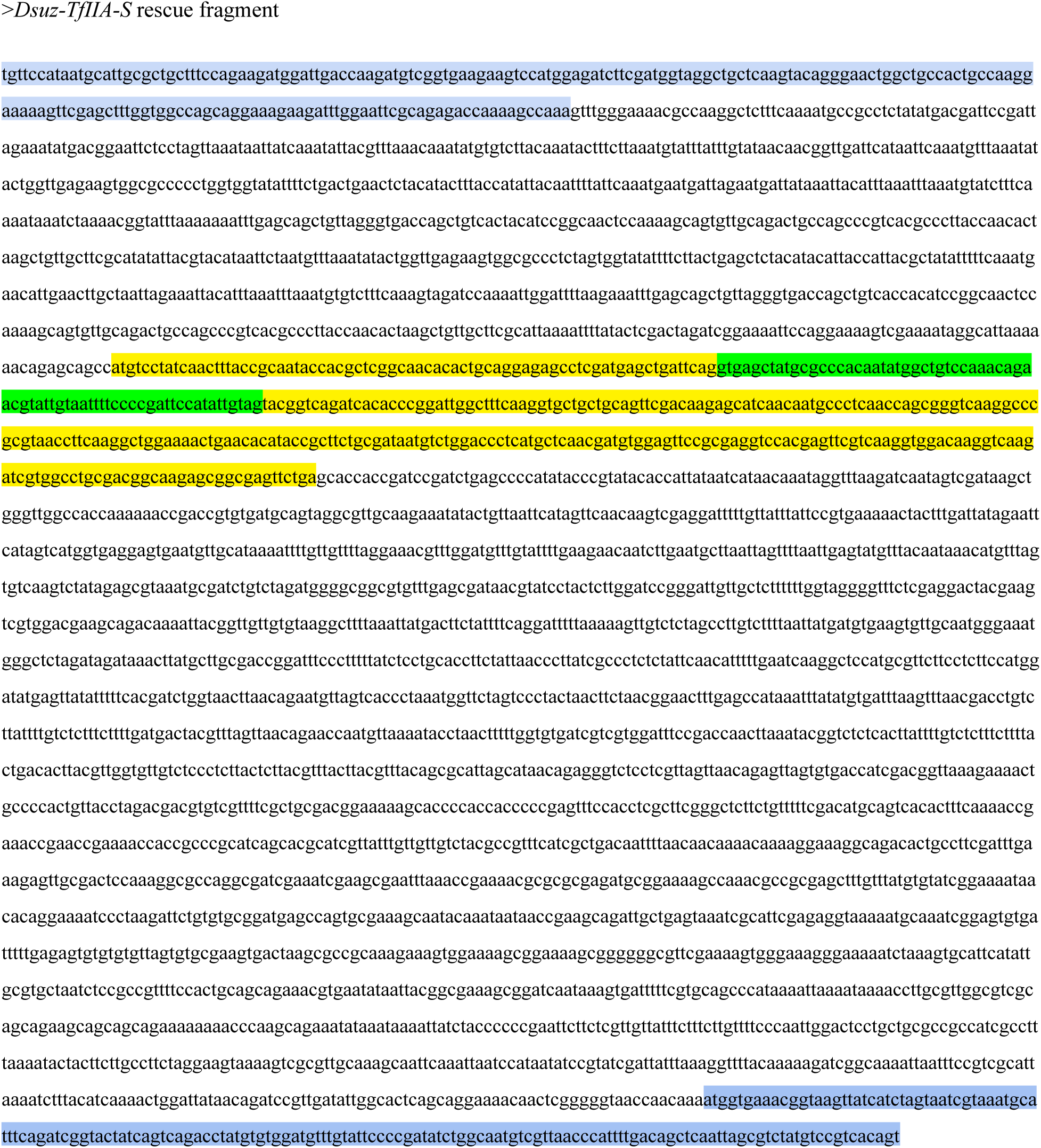
*Dsuz-TfIIA-S* rescue fragment fasta file. Exons in yellow, intron in green, up- and downstream annotated gene neighbors in blue, total length=3802 bp. The *Rescue* fragment was chosen to include sequence from neighboring genes to maximize the likelihood that all regulatory sequences needed for normal expression of the *Rescue* transgene are present.

**Fig S8.**
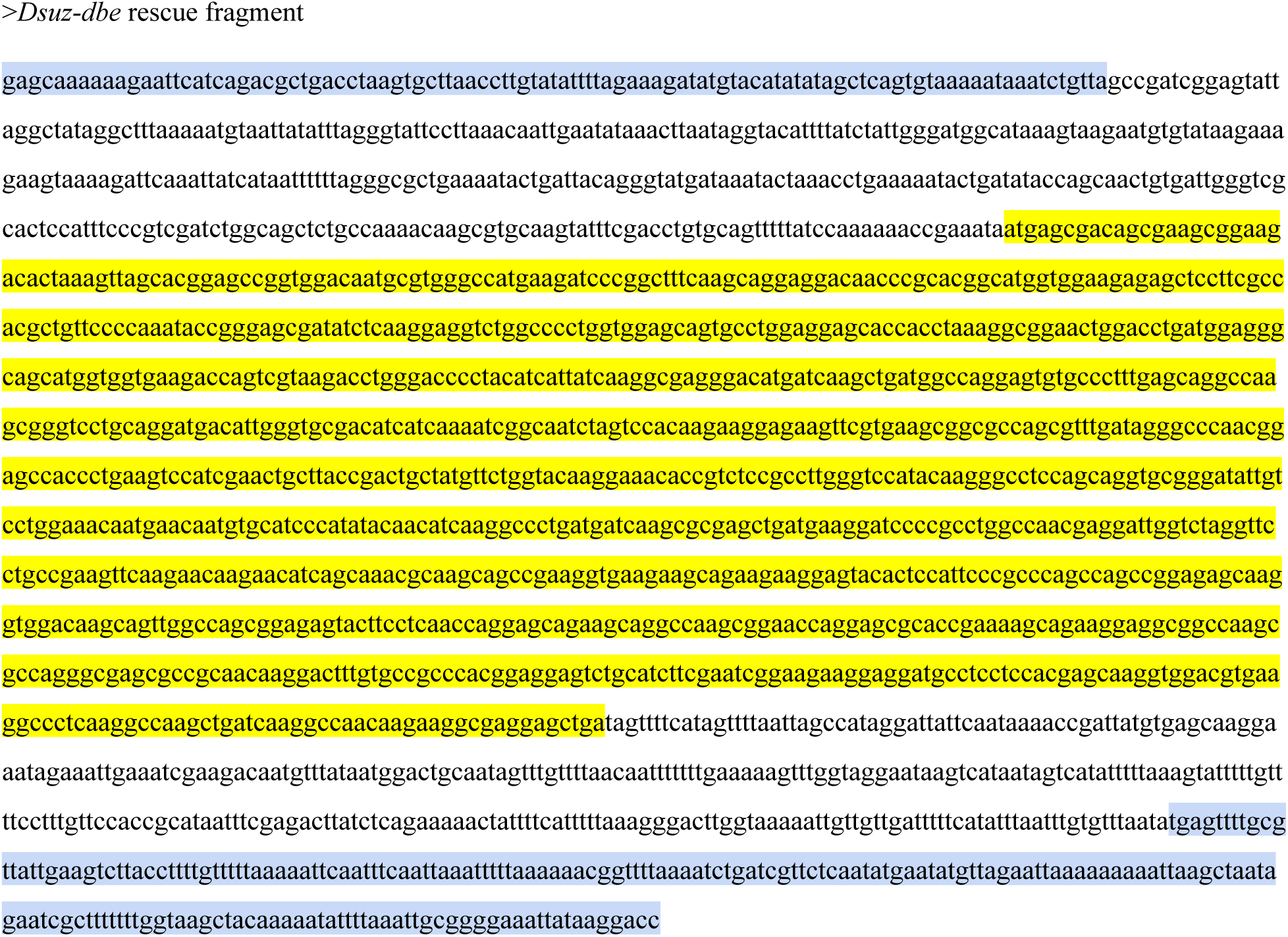
*Dsuz-dbe* rescue fragment fasta file. Exon in yellow, up- and downstream annotated gene neighbors in blue, total length=1961 bp. The *Rescue* fragment was chosen to include sequence from neighboring genes to maximize the likelihood that all regulatory sequences needed for normal expression of the *Rescue* transgene are present.

### Supplementary Tables

**Table S1:**
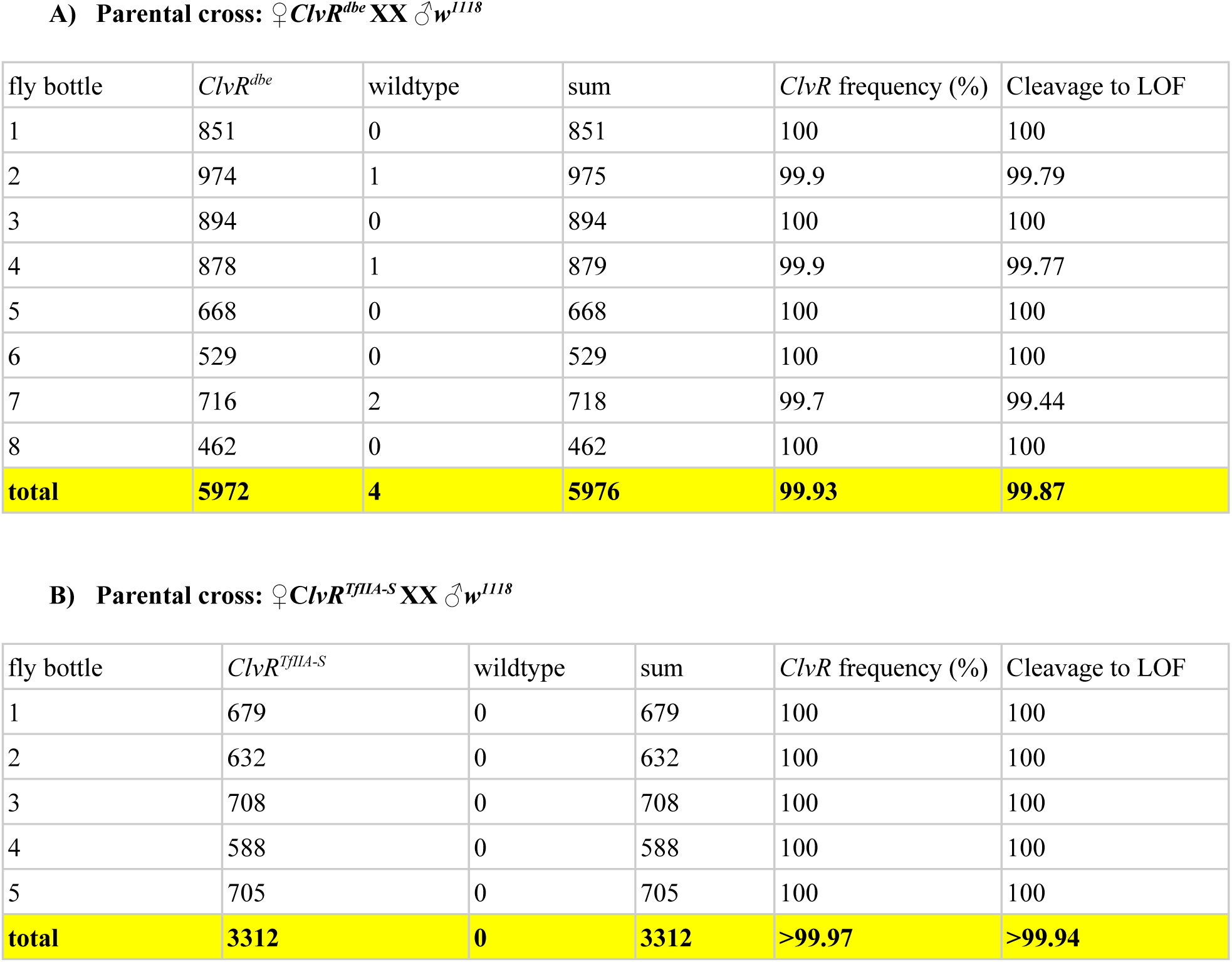
Rates of LOF allele creation in the combined maternal germline and zygote of ClvR^dbe^ and ClvR^TfIIA-S^. We scored the genotype of offspring from a cross of *ClvR*/+ heterozygous mothers to wildtype *w^1118^* males. The genotype frequencies of the offspring of these crosses were used to calculate a combined LOF allele creation rate coming from cleavage in the germline and cleavage of the paternal target gene allele due to maternal carry over. Crosses were set up in bottles with ∼40 *ClvR*/+ virgins each. The 4 presumably wildtype flies (escapers) were analyzed further (see Table S4)

**Table S2:** Rates of LOF allele creation in the paternal germline. The target genes of our *ClvR* lines are recessive lethal. To determine rates of cleavage and LOF allele creation in progeny from heterozygous *ClvR*/+ males we crossed them to deficiency stocks that completely lacked the target gene locus. The deficiency-bearing chromosome was maintained in trans to a balancer chromosome that is dominantly marked and that is wildtype for the target essential gene. By focusing on the offspring that carry the deficiency we can calculate the rate at which LOF alleles are created in the male germline by dividing the number of flies that carry the deficiency and *ClvR* by half the number of flies that carry the deficiency: male germline LOF allele rate= [*Df(dbe),ClvR^dbe^*] / sum[*Df(dbe)/2*]. The 21 flies that carried the deficiency but not *ClvR* (escapers) were analyzed further below, Table S5)

| vial | *Df(dbe), ClvR<sup>dbe</sup>* | *Df(dbe)* | *CyO, ClvR<sup>dbe</sup>* | *CyO* | sum *Df(dbe)* | *ClvR* frequency (%) | Cleavage to LOF |
| --- | --- | --- | --- | --- | --- | --- | --- |
| 1 | 15 | 0 | 21 | 9 | 15 | 100 | 100 |
| 2 | 29 | 0 | 25 | 29 | 29 | 100 | 100 |
| 3 | 9 | 0 | 3 | 5 | 9 | 100 | 100 |
| 4 | 43 | 4 | 50 | 30 | 47 | 91.5 | 82.98 |
| 5 | 16 | 0 | 15 | 16 | 16 | 100 | 100 |
| 6 | 5 | 0 | 7 | 2 | 5 | 100 | 100 |
| 7 | 18 | 4 | 25 | 25 | 22 | 81.8 | 63.64 |
| 8 | 23 | 0 | 26 | 25 | 23 | 100 | 100 |
| 9 | 31 | 0 | 35 | 28 | 31 | 100 | 100 |
| 10 | 17 | 0 | 28 | 26 | 17 | 100 | 100 |
| 11 | 14 | 0 | 19 | 29 | 14 | 100 | 100 |
| 12 | 34 | 2 | 38 | 33 | 36 | 94.4 | 88.89 |
| 13 | 15 | 0 | 17 | 14 | 15 | 100 | 100 |
| 14 | 40 | 1 | 44 | 31 | 41 | 97.6 | 95.12 |
| 15 | 22 | 2 | 24 | 23 | 24 | 91.7 | 83.33 |
| 16 | 33 | 0 | 28 | 28 | 33 | 100 | 100 |
| 17 | 33 | 0 | 25 | 15 | 33 | 100 | 100 |
| 18 | 21 | 0 | 21 | 27 | 21 | 100 | 100 |
| 19 | 43 | 4 | 36 | 42 | 47 | 91.5 | 82.98 |
| 20 | 24 | 0 | 18 | 26 | 24 | 100 | 100 |
| 21 | 18 | 0 | 27 | 23 | 18 | 100 | 100 |
| 22 | 23 | 0 | 22 | 23 | 23 | 100 | 100 |
| 23 | 26 | 1 | 27 | 35 | 27 | 96.3 | 92.60 |
| 24 | 31 | 0 | 35 | 40 | 31 | 100 | 100 |
| 25 | 19 | 0 | 13 | 25 | 19 | 100 | 100 |
| 26 | 19 | 0 | 19 | 18 | 19 | 100 | 100 |
| 27 | 27 | 0 | 20 | 26 | 27 | 100 | 100 |
| 28 | 20 | 2 | 32 | 33 | 22 | 90.9 | 81.82 |
| 29 | 12 | 0 | 18 | 18 | 12 | 100 | 100 |
| 30 | 29 | 0 | 22 | 35 | 29 | 100 | 100 |
| 31 | 29 | 1 | 38 | 39 | 30 | 96.7 | 93.33 |
| 32 | 20 | 0 | 20 | 21 | 20 | 100 | 100 |
| 33 | 18 | 0 | 20 | 19 | 18 | 100 | 100 |
| **total** | **776** | **21** | **818** | **818** | **797** | **97.4** | **94.73** |

**B) Parental cross: ♂ *ClvR<sup>TfIIA-S</sup>* XX ♀ *Df(TfIIA-S)/TM6B,Tb***
| vial | *Df(TfIIA-S)/ClvR<sup>TfIIA-S</sup>* | *Df(TfIIA-S)* | *ClvR<sup>TfIIA-S</sup>/TM6B,*<br>*Tb* | *TM6B,Tb* | sum<br>*Df(TfIIA-S)* | *ClvR*<br>frequency<br>(%) | Cleavage to LOF |
| --- | --- | --- | --- | --- | --- | --- | --- |
| 1 | 32 | 0 | 27 | 26 | 32 | 100 | 100 |
| 2 | 20 | 0 | 28 | 21 | 20 | 100 | 100 |
| 3 | 26 | 1 | 28 | 32 | 27 | 96.3 | 92.60 |
| 4 | 34 | 0 | 36 | 23 | 34 | 100 | 100 |
| 5 | 27 | 0 | 28 | 26 | 27 | 100 | 100 |
| 6 | 34 | 0 | 29 | 24 | 34 | 100 | 100 |
| 7 | 29 | 0 | 35 | 35 | 29 | 100 | 100 |
| 8 | 13 | 0 | 36 | 27 | 13 | 100 | 100 |
| 9 | 19 | 0 | 16 | 15 | 19 | 100 | 100 |
| 10 | 23 | 0 | 10 | 13 | 23 | 100 | 100 |
| 11 | 28 | 0 | 27 | 24 | 28 | 100 | 100 |
| 12 | 3 | 0 | 2 | 3 | 3 | 100 | 100 |
| 13 | 32 | 0 | 42 | 37 | 32 | 100 | 100 |
| 14 | 26 | 0 | 20 | 18 | 26 | 100 | 100 |
| 15 | 36 | 0 | 41 | 31 | 36 | 100 | 100 |
| 16 | 19 | 0 | 17 | 19 | 19 | 100 | 100 |
| 17 | 26 | 0 | 32 | 31 | 26 | 100 | 100 |
| 18 | 11 | 0 | 13 | 10 | 11 | 100 | 100 |
| 19 | 24 | 0 | 31 | 30 | 24 | 100 | 100 |
| 20 | 35 | 0 | 42 | 31 | 35 | 100 | 100 |
| 21 | 30 | 0 | 31 | 34 | 30 | 100 | 100 |
| 22 | 11 | 1 | 21 | 18 | 12 | 91.7 | 83.33 |
| 23 | 28 | 0 | 24 | 27 | 28 | 100 | 100 |
| 24 | 34 | 0 | 32 | 28 | 34 | 100 | 100 |
| 25 | 23 | 0 | 23 | 24 | 23 | 100 | 100 |
| 26 | 31 | 0 | 27 | 22 | 31 | 100 | 100 |
| 27 | 18 | 0 | 25 | 24 | 18 | 100 | 100 |
| total | 672 | 2 | 723 | 653 | 674 | 99.7 | 99.41 |

**Table S3:** Rates of LOF allele creation in the combined maternal germline and zygote of the second generation *ClvR* elements *ClvR^dbe^+R^tko^* and *ClvR^tko^+R^dbe^*. We scored the genotype of offspring from a cross of (*ClvR^n+1^+R^n^*)/+ heterozygous mothers to wildtype *w^1118^* males. The genotype frequencies of the offspring of these crosses were used to calculate a combined LOF allele rate coming from cleavage in the germline and cleavage of the paternal target gene allele due to maternal carry over. Crosses were set up in bottles with ∼40 (*ClvR^n+1^+R^n^*)/+ virgins each.

| fly bottle | *ClvR<sup>dbe</sup>+R<sup>tko</sup>* | wildtype | sum | *ClvR* frequency (%) | Cleavage to LOF |
| --- | --- | --- | --- | --- | --- |
| 1 | 1138 | 0 | 1138 | 100 | 100 |
| 2 | 1053 | 0 | 1053 | 100 | 100 |
| 3 | 812 | 0 | 812 | 100 | 100 |
| **total** | **3003** | **0** | **3003** | **>99.97** | **>99.87** |

**B) Parental cross: ♀(*ClvR<sup>tko</sup>+R<sup>dbe</sup>*)/+ XX ♂*w<sup>1118</sup>***
| fly bottle | *ClvR<sup>tko</sup>+R<sup>dbe</sup>* | wildtype | sum | *ClvR* frequency (%) | Cleavage to LOF |
| --- | --- | --- | --- | --- | --- |
| 1 | 1106 | 0 | 1106 | 100 | 100 |
| 2 | 1329 | 0 | 1329 | 100 | 100 |
| 3 | 1164 | 0 | 1164 | 100 | 100 |
| **total** | **3599** | **0** | **3599** | **>99.97** | **>99.89%** |

**Table S4:** Analysis of escapers from crosses of heterozygous ClvR^dbe^/+ females to w^1118^ in Table S1. Out of 5972 flies scored we found 4 flies that did not carry the *ClvR* marker. We crossed all 4 of them again to heterozygous *ClvR*/+ females to test whether the escaper chromosomes remained sensitive to *ClvR*. Results of this cross are in the last 3 columns of the table below. After allowing the escaper flies to mate we extracted genomic DNA and sequenced over the gRNA target sites. Sequencing results are shown in the table below. ‘+’ indicates an unaltered target site, del indicates a deletion followed by the number of deleted bp. One fly died for which we couldn’t obtain a sequencing signal. The other three had a common 3bp in frame deletion at the target site for gRNA4. Two of these flies had an additional 3bp in frame deletion at target site for gRNA3. Target sites for gRNA1 and gRNA2 were unaltered. Backcrosses of these flies to *ClvR*/+ females showed that the escaper chromosome could still be cleaved/mutated (Cleavage to LOF of 93.75 and 100%, 1 fly was sterile).

| Fly ID | gRNA1 | gRNA2 | gRNA3 | gRNA4 | tomato+ | tomato- | *ClvR* frequency (%) | Cleavage to LOF |
| --- | --- | --- | --- | --- | --- | --- | --- | --- |
| 2.1 | + | + | 3bp del | 3bp del | 62 | 2 | 96.88 | 93.75 |
| 4.1 | no signal | + | 3bp del | 3bp del | sterile | sterile | NA | NA |
| 6.1 | + | + | + | 3bp del | 51 | 0 | 100 | 100 |

**Table S5:** Analysis of escapers from crosses of heterozygous *Clvr^dbe^*/+ and *ClvR/^TfIIA-S^*/+ males to the deficiency strains from Table S2. For the cross of **♂***ClvR^dbe^* XX *Df(dbe)/CyO* we found 21 escapers among 2433 flies scored. For the cross of **♂***ClvR^TfIIA-S^* XX *Df(TfIIA-S)/TM3,Tb* we found 2 escapers among 2050 flies scored. We extracted genomic DNA from all of the escapers and sequenced over the target region. In addition, male escapers were backcrossed to heterozygous *ClvR*-bearing females to check whether the escaper chromosome could still be cleaved and mutated to LOF. Results are summarized in the table below. ‘+’ stands for unaltered target site, SNP is likely a pre-existing polymorphism in the target site, ‘del’ stands for deletion followed by the number of deleted bases, ‘sub’ stands for substitution of bases. The last four columns show the results of the backcrosses of male escapers to heterozygous *ClvR/+* females to check whether the escaped target locus could still be cleaved. For the 21 escapers coming from male *ClvR^dbe^/+* fathers, we found a 3bp in frame deletion at the target site for gRNA4 in five flies. One fly had a polymorphism at the target site of gRNA3 and one fly had a 2bp substitution at target site of gRNA3. All other target sites were unaltered. 12 of the 21 flies were males, which were backcrossed to *ClvR/+* females. In one of theses crosses the cleavage rate to LOF was 85%, for all the others it was 100% (see Table S5A), again showing that the escaper chromosomes were not resistant to Cas9 cleavage. The 2 escapers coming from *ClvR^TfIIA-S^* fathers had a polymorphism at the target site of gRNA2, while all the other target sites remained unaltered. When the 2 escapers were backcrossed to *ClvR*/+ females, the cleavage rate to LOF in the progeny was 100% (Table S5B).

| Fly ID | gRNA1 | gRNA2 | gRNA3 | gRNA4 | tomato+ | tomato- | *ClvR*<br>frequency(%) | Cleavage to<br>LOF | notes |
| --- | --- | --- | --- | --- | --- | --- | --- | --- | --- |
| 4.1 | + | + | + | + | 74 | 0 | 100 | 100 | male |
| 4.2 | + | + | + | + | 64 | 0 | 100 | 100 | male |
| 4.3 | + | + | + | 3bp del | NA | NA | NA | NA | female |
| 4.4 | + | + | + | + | 71 | 0 | 100 | 100 | male |
| 7.1 | + | + | SNP | 3bp del | NA | NA | NA | NA | female |
| 7.2 | + | + | + | + | 76 | 0 | 100 | 100 | male |
| 7.3 | + | + | + | + | 53 | 4 | 93.0 | 84.9 | male |
| 7.4 | + | + | + | + | NA | NA | NA | NA | female |
| 12.1 | + | + | + | + | NA | NA | NA | NA | female |
| 12.2 | + | + | + | + | 66 | 0 | 100 | 100 | male |
| 14.1 | + | + | + | 3bp del | NA | NA | NA | NA | female |
| 15.1 | + | + | + | + | sterile | sterile | NA | NA | male |
| 15.2 | + | + | + | + | 54 | 0 | 100 | 100 | male |
| 19.1 | + | + | 2bp sub | + | NA | NA | NA | NA | female |
| 19.2 | + | + | + | + | NA | NA | NA | NA | female |
| 19.3 | + | + | + | 3bp del | 77 | 0 | 100 | 100 | male |
| 19.4 | + | + | + | 3bp del | 71 | 0 | 100 | 100 | male |
| 23.1 | + | + | + | + | NA | NA | NA | NA | female |
| 28.1 | + | + | + | + | 68 | 0 | 100 | 100 | male |
| 28.2 | + | + | + | + | 61 | 0 | 100 | 100 | male |
| 31.1 | + | + | + | + | 66 | 0 | 100 | 100 | male |

| Fly ID | gRNA<br>1 | gRNA2 | gRNA3 | gRNA4 | tomato+ | tomato- | *ClvR* frequency<br>(%) | Cleavage to<br>LOF | notes |
| --- | --- | --- | --- | --- | --- | --- | --- | --- | --- |
| 3.1 | + | SNP | + | + | 71 | 0 | 100 | 100 | male |
| 22.1 | + | SNP | + | + | 89 | 0 | 100 | 100 | male |

**Table S6:** Cleavage events in chromosomes exposed to ClvR^dbe^ for the first time. We sequenced the target locus of *ClvR^dbe^* flies in a chromosome that was exposed to *ClvR^dbe^* for the first time. The crossing scheme to obtain these flies is provided below. We started with the cross to determine the male germline cleavage rate to LOF by crossing ♂*ClvR^dbe^* XX ♀*Df(dbe)/CyO* (Data in Table S2). This cross produced ♀*dbe^-^/CyO;ClvR^dbe^.* The *CyO* balancer chromosome came from the deficiency stock and is now exposed to the *ClvR* element for the first time. Next we backcrossed these females again to males of the deficiency stock: ♀*dbe^-^/CyO;ClvR^dbe^* XX ♂*Df^dbe^/CyO*. Among the progeny of this cross were flies that had the *CyO* balancer chromosome of the mother in trans to the deficiency from the father. Flies with that genotype were used to sequence over the target region on the *CyO* balancer chromosome. To simplify the scheme below it is assumed that *ClvR^dbe^* always creates mutations at the wildtype *dbe* locus, indicated as *dbe^-^*.

Sequencing results of *Df(dbe)/CyO;ClvR<sup>dbe</sup>* flies:
| Fly ID | gRNA1 | gRNA2 | gRNA3 | gRNA4 |
| --- | --- | --- | --- | --- |
| 27 | + | SNP | del between gRNA3 and 4 |  |
| 28 | + | SNP | 2bp indel | 37bp del |
| 29 | + | SNP | 9bp del | 8bp del |
| 30 | + | SNP | 11bp insertion | 6bp del |
| 31 | + | SNP | + | + |
| 32 | del between gRNA1 and 2-3 |  | 9bp del | 3bp del |
| 33 | 17bp del | 6bp del | 4bp del | 4bp indel |
| 34 | + | SNP | del between gRNA3 and 4 |  |
| 35 | 3bp del | 5bp indel | 11bp insertion | 3bp del |
| 36 | + | SNP | del between gRNA3 and 4 |  |
| 37 | + | SNP | + | + |
| 38 | + | SNP | 11bp insertion | 3bp del |
| 39 | + | SNP | 11bp indel | 38 bp del |
| 40 | + | SNP | 3 bp indel | 7bp del |
| 41 | + | SNP | 13 bp deletion | 5bp insertion |
| 42 | + | SNP | 4 bp deletion | 2bp deletion |

**Table S7:**
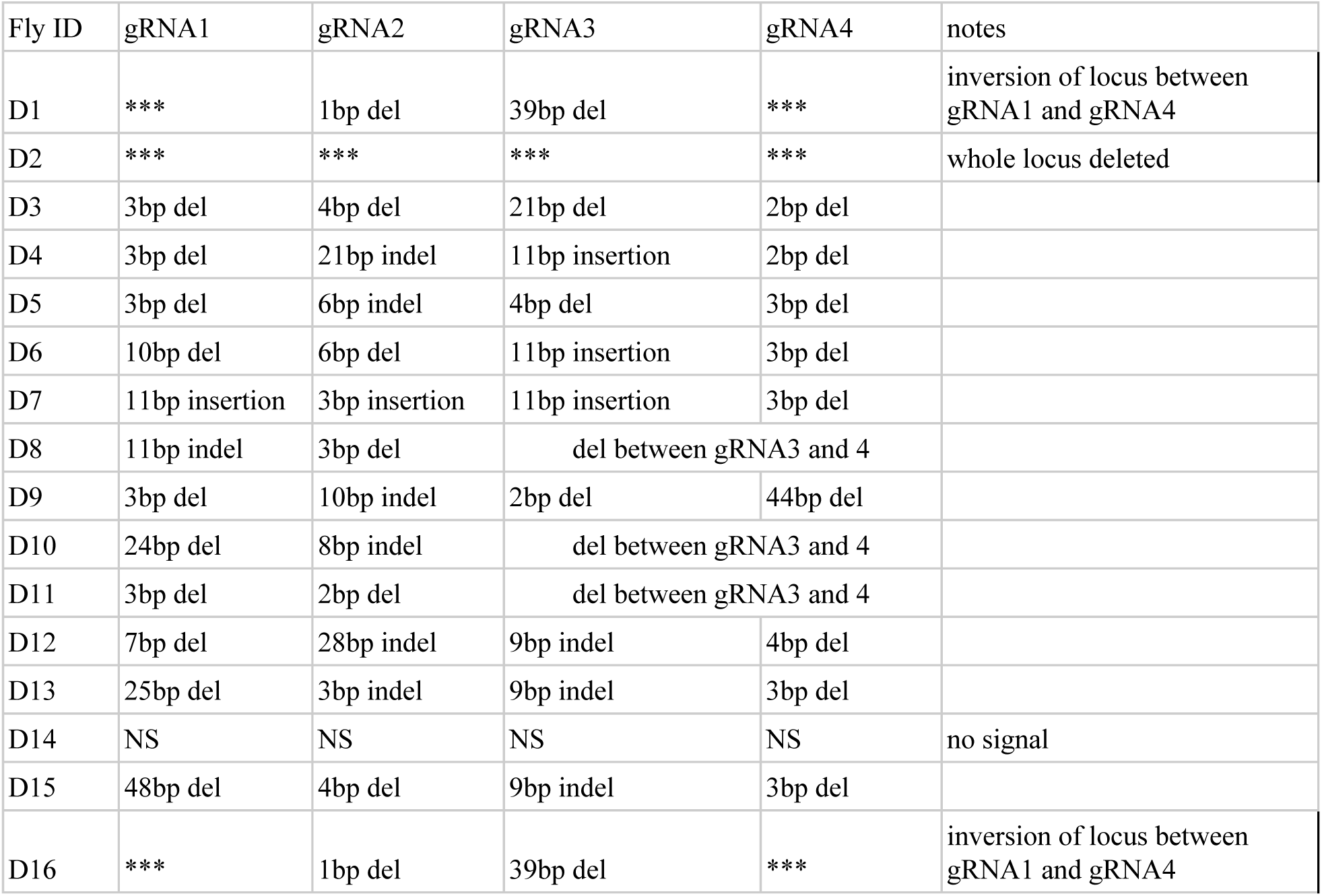
Cleavage events in chromosomes of *ClvR^dbe^* drive populations after 22 Generations. We took 4 flies of each drive replicate bottle, crossed them individually to the deficiency stock, and sequenced progeny of this cross that had a presumably cleaved/mutated *dbe* locus in trans to the deficiency. Sequencing results are summarized in the table below.

**Table S8:** Cleavage events in chromosomes exposed to ClvR^TfIIA-S^. For *ClvR^TfIIA-S^*, the *ClvR* selfish element as well as the target gene are located on the same chromosome. Therefore, we could not set up crosses easily where it would be clear that the cleaved target locus was only generated in that generation. However, as an attempt to generate what were likely new mutations at the *TfIIA-S* locus we took heterozygous *ClvR/+* females and outcrossed them to *w^1118^* males for 5 generations. We then crossed *ClvR/+* females to a deficiency strain for the *TfIIA-S* locus and sequenced the target locus in progeny that were ClvR*^TfIIA-S^*/*Df*. Results are summarized in the following table.

| Fly ID | gRNA1 | gRNA2 | gRNA3 | gRNA4 | notes |
| --- | --- | --- | --- | --- | --- |
| 1 | 3bp del | 2bp ins, 3 bp sub | 19bp del | 8bp del |  |
| 2 | del between gRNA1 and gRNA2 |  | 19 bp del | 26bp del |  |
| 3 | 9bp del | 2bp del | 11bp del | 8bp del |  |
| 4 | deletion between gRNA1 and gRNA4 |  |  |  |  |
| 5 | 3bp del | 2bp ins, 3 bp sub | 19bp del | 8bp del |  |
| 6 | del between gRNA1 and gRNA2 |  | 19 bp del | 26bp del |  |
| 7 | del between gRNA1 and gRNA2 |  | 19 bp del | 26bp del |  |
| 8 | del between gRNA1 and gRNA2 |  | 19 bp del | 26bp del |  |
| 9 | del between gRNA1 and gRNA2 |  | 19 bp del | 26bp del |  |
| 10 | del between gRNA1 and gRNA2 |  | 19 bp del | 26bp del |  |
| 11 | del between gRNA1 and gRNA2 |  | 19 bp del | 26bp del |  |
| 12 | 9bp del | 2bp del | 11bp del | 8bp del |  |
| 13 | NS | NS | NS | NS | no signal |
| 14 | del between gRNA1 and gRNA2 |  | 19 bp del | 26bp del |  |
| 15 | del between gRNA1 and gRNA2 |  | 19 bp del | 26bp del |  |
| 16 | del between gRNA1 and gRNA2 |  | 19 bp del | 26bp del |  |

**Table S9:** Cleavage events in chromosomes of *ClvR^TfIIA-S^* drive populations after 22 Generations. We took 4 flies from each drive replicate bottle, crossed them individually to the *Df* stock for the *TfIIA-S* region, and sequenced progeny of this cross that were *ClvR^TfIIA-S^* and carried the *Df* chromosome. These flies presumably had a cleaved *TfIIA-S* locus in trans to the deficiency. Sequencing results are summarized in the table below.

| Fly ID | gRNA1 | gRNA2 | gRNA3 | gRNA4 | notes |
| --- | --- | --- | --- | --- | --- |
| T1 | 3bp del | 6bp del | 11bp del | 30bp del |  |
| T2 | del between gRNA1 and 2 |  | 19bp del | 26 bp del |  |
| T3 | del between gRNA1 and 2 |  | 19bp del | 26 bp del |  |
| T4 | del between gRNA1 and 2 |  | 19bp del | 26 bp del |  |
| T5 | del between gRNA1 and 2 |  | 19bp del | 26 bp del |  |
| T6 | del between gRNA1 and 2 |  | 19bp del | 26 bp del |  |
| T7 | del between gRNA1 and 2 |  | 19bp del | 26 bp del |  |
| T8 | del between gRNA1 and 2 |  | 19bp del | 26 bp del |  |
| T9 | NS | NS | NS | NS | no signal |
| T10 | del between gRNA1 and 2 |  | 19bp del | 26 bp del |  |
| T11 | del between gRNA1 and 2 |  | 19bp del | 26 bp del |  |
| T12 | del between gRNA1 and 2 |  | 19bp del | 26 bp del |  |
| T13 | del between 1.1kb upstream of gRNA1 and 3 |  |  | 3bp del |  |
| T14 | del between gRNA1 and 2 |  | 19bp del | 26 bp del |  |
| T15 | del between gRNA1 and 2 |  | 19bp del | 4bp del |  |
| T16 | 2bp del | 1bp del | del between gRNA3 and 4 |  |  |

**Table S10:** Allele frequency of ClvR elements in gene drive experiments. The number of *ClvR* homozygotes and heterozygotes in different gene drive populations was determined by outcrossing 100 males from each gene drive bottle to *w^1118^* females. If 100% of the progeny of these crosses carried the *ClvR^n^* marker, the male was homozygous for *ClvR^n^*. If half carried the *ClvR^n^* marker the father was heterozygous. In addition, virgins were collected from the bottles at the assayed generations and used to seed the *ClvR^n^+R^n-1^* drive experiments presented in Fig. 4.

| *ClvR* target | Replicate | Generation | *ClvR<sup>n</sup>* allele frequency [%] |
| --- | --- | --- | --- |
| *dbe* | A | 22 | 90.43 |
| *dbe* | B | 22 | 82.76 |
| *dbe* | C | 22 | 87.36 |
| *dbe* | D | 22 | 89.20 |
| *TfIIA-S* | A | 22 | 91.07 |
| *TfIIA-S* | B | 22 | 93.75 |
| *TfIIA-S* | C | 22 | 94.71 |
| *TfIIA-S* | D | 22 | 95.70 |
| *tko* | A | 32 | 93.96 |
| *tko* | B | 32 | 95.60 |
| *tko* | C | 32 | 97.65 |
| *tko* | D | 32 | 100.00 |
| *tko* | E | 32 | 100.00 |

**Table S11.**
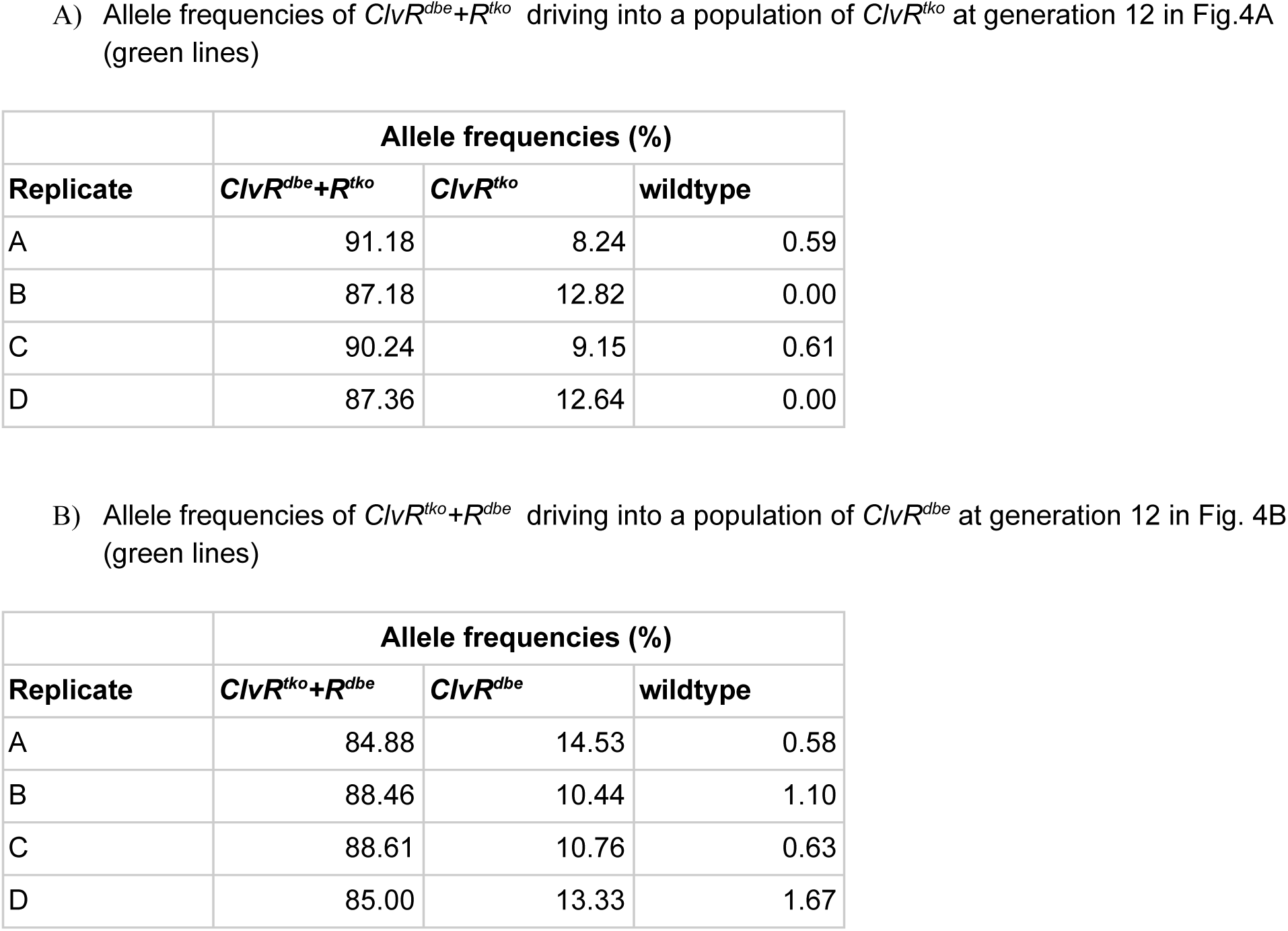
Allele frequencies of *ClvR^n+1^+R^n^, ClvR^n^,* and wildtype in 2nd generation *ClvR* gene drive experiments at generation 12. The allele frequencies in these gene drive populations was determined by outcrossing 100 males from each gene drive bottle to *w^1118^* females. If 100% of the progeny of these crosses carried the *ClvR^n+1^+R^n^* marker, the male was homozygous for *ClvR^n+1^+R^n^*. If half carried the *ClvR^n^* marker but not *ClvR^n+1^+R^n^* the father was transheterozygous for the 2nd and 1st generation *ClvR* element. If half carried no marker the father was heterozygous. Starting allele frequency for *ClvR^n+1^+R^n^* at generation 0 in all gene drive experiments was 25% (starting allele frequencies of *ClvR^n^* are in Table S10). Note how *ClvR^n+1^+R^n^* increases in frequency at the cost of *ClvR^n^*. Phenotype frequencies are plotted in Fig. 4 (green lines).

**Table S12.** Allele frequencies of ClvR^n+1^+R^n^ driving into populations of w^1118^ at generation 12. The allele frequencies in these gene drive populations was determined by outcrossing 100 males from each gene drive bottle to *w^1118^* females. If 100% of the progeny carried the *ClvR^n+1^+R^n^* marker, the male was homozygous for *ClvR^n+1^+R^n^*. If half carried no marker the father was heterozygous. Starting allele frequency for *ClvR^n+1^+R^n^* at generation 0 in all gene drive experiments was 25%. Note how *ClvR^n+1^+R^n^* increases in frequency at the cost wildtype (*w^1118^*). Phenotype frequencies are plotted in Fig. 4 (red lines)

|  | Allele frequencies (%) |  |
| --- | --- | --- |
| Replicate | $ClvR^{dbe}+R^{tko}$ | wildtype |
| A | 84.76 | 15.24 |
| B | 81.33 | 18.67 |
| C | 84.83 | 15.17 |
| D | 87.34 | 12.66 |

|  | Allele frequencies (%) |  |
| --- | --- | --- |
| Replicate | $ClvR^{tko}+R^{dbe}$ | wildtype |
| A | 83.73 | 16.27 |
| B | 87.67 | 12.33 |
| C | 82.72 | 17.28 |
| D | 86.14 | 13.86 |

## References

1. P. Schmid-Hempel, Evolutionary ecology of insect immune defenses. Annu. Rev. Entomol. 50, 529–551 (2005).

2. F. Tripet, F. Aboagye-Antwi, H. Hurd, Ecological immunology of mosquito-malaria interactions. Trends Parasitol. 24, 219–227 (2008).

3. J. R. Carter, et al., Suppression of the Arboviruses Dengue and Chikungunya Using a Dual-Acting Group-I Intron Coupled with Conditional Expression of the Bax C-Terminal Domain. PLoS One 10, e0139899 (2015).

4. X. J. Gao, L. S. Chong, M. S. Kim, M. B. Elowitz, Programmable protein circuits in living cells. Science 361, 1252–1258 (2018).

5. H. R. Braig, G. Yan, “The spread of genetic constructs in natural insect populations” in Genetically Engineered Organisms: Assessing Environmental and Human Health Effects, D. K. Letourneau, B. E. Burrows, Eds. (CRC Press, 2001), pp. 251–314.

6. S. Davis, N. Bax, P. Grewe, Engineered underdominance allows efficient and economical introgression of traits into pest populations. J. Theor. Biol. 212, 83–98 (2001).

7. A. Burt, Site-specific selfish genes as tools for the control and genetic engineering of natural populations. Proc. Biol. Sci. 270, 921–928 (2003).

8. F. Gould, P. Schliekelman, Population genetics of autocidal control and strain replacement. Annu. Rev. Entomol. 49, 193–217 (2004).

9. C.-H. Chen, et al., A synthetic maternal-effect selfish genetic element drives population replacement in Drosophila. Science 316, 597–600 (2007).

10. P. M. Altrock, A. Traulsen, R. G. Reeves, F. A. Reed, Using underdominance to bi-stably transform local populations. J. Theor. Biol. 267, 62–75 (2010).

11. J. M. Marshall, B. A. Hay, Inverse Medea as a novel gene drive system for local population replacement: a theoretical analysis. J. Hered. 102, 336–341 (2011).

12. J. M. Marshall, G. W. Pittman, A. B. Buchman, B. A. Hay, Semele: a killer-male, rescue-female system for suppression and replacement of insect disease vector populations. Genetics 187, 535–551 (2011).

13. P. M. Altrock, A. Traulsen, F. A. Reed, Stability properties of underdominance in finite subdivided populations. PLoS Comput. Biol. 7, e1002260 (2011).

14. J. M. Marshall, B. A. Hay, General principles of single-construct chromosomal gene drive. Evolution 66, 2150–2166 (2012).

15. O. S. Akbari, et al., A synthetic gene drive system for local, reversible modification and suppression of insect populations. Curr. Biol. 23, 671–677 (2013).

16. K. M. Esvelt, A. L. Smidler, F. Catteruccia, G. M. Church, Concerning RNA-guided gene drives for the alteration of wild populations. Elife 3 (2014).

17. C. S. Gokhale, R. G. Reeves, F. A. Reed, Dynamics of a combined Medea-underdominant population transformation system. BMC Evol. Biol. 14, 98 (2014).

18. R. G. Reeves, J. Bryk, P. M. Altrock, J. A. Denton, F. A. Reed, First steps towards underdominant genetic transformation of insect populations. PLoS One 9, e97557 (2014).

19. J. Sudweeks, et al., Locally Fixed Alleles: A method to localize gene drive to island populations. Sci. Rep. 9, 15821 (2019).

20. A. Nash, et al., Integral Gene Drives: an “operating system” for population replacement. bioRxiv, 356998 (2018).

21. V. L. Del Amo, et al., Split-gene drive system provides flexible application for safe laboratory investigation and potential field deployment. bioRxiv, 684597 (2019).

22. J. Champer, et al., A toxin-antidote CRISPR gene drive system for regional population modification. bioRxiv, 628354 (2019).

23. O. S. Akbari, et al., Novel Synthetic Medea Selfish Genetic Elements Drive Population Replacement in Drosophila; a Theoretical Exploration of Medea-Dependent Population Suppression. ACS Synth. Biol. (2012) https://doi.org/10.1021/sb300079h.

24. A. B. Buchman, T. Ivy, J. M. Marshall, O. S. Akbari, B. A. Hay, Engineered Reciprocal Chromosome Translocations Drive High Threshold, Reversible Population Replacement in Drosophila. ACS Synth. Biol. (2018) https://doi.org/10.1021/acssynbio.7b00451.

25. G. Oberhofer, T. Ivy, B. A. Hay, Cleave and Rescue, a novel selfish genetic element and general strategy for gene drive. Proc. Natl. Acad. Sci. U. S. A. (2019) https://doi.org/10.1073/pnas.1816928116.

26. K. Kyrou, et al., A CRISPR-Cas9 gene drive targeting doublesex causes complete population suppression in caged Anopheles gambiae mosquitoes. Nat. Biotechnol. (2018) https://doi.org/10.1038/nbt.4245.

27. J. Champer, et al., Reducing resistance allele formation in CRISPR gene drive. Proc. Natl. Acad. Sci. U. S. A. 115, 5522–5527 (2018).

28. G. Oberhofer, T. Ivy, B. A. Hay, Behavior of homing endonuclease gene drives targeting genes required for viability or female fertility with multiplexed guide RNAs. Proc. Natl. Acad. Sci. U. S. A. (2018) https://doi.org/10.1073/pnas.1805278115.

29. B. A. Hay, et al., Engineering the genomes of wild insect populations: challenges, and opportunities provided by synthetic Medea selfish genetic elements. J. Insect Physiol. 56, 1402–1413 (2010).

30. J. Champer, I. Kim, S. E. Champer, A. G. Clark, P. W. Messer, Performance analysis of novel toxin-antidote CRISPR gene drive systems. bioRxiv, 628362 (2019).

31. H. Y. Chan, S. Brogna, C. J. O’Kane, Dribble, the Drosophila KRR1p homologue, is involved in rRNA processing. Mol. Biol. Cell 12, 1409–1419 (2001).

32. K. Yokomori, et al., Drosophila TFIIA directs cooperative DNA binding with TBP and mediates transcriptional activation. Genes Dev. 8, 2313–2323 (1994).

33. K. A. Molla, Y. Yang, CRISPR/Cas-Mediated Base Editing: Technical Considerations and Practical Applications. Trends Biotechnol. 37, 1121–1142 (2019).

34. A. V. Anzalone, et al., Search-and-replace genome editing without double-strand breaks or donor DNA. Nature (2019) https://doi.org/10.1038/s41586-019-1711-4.

35. F. Lansing, et al., A heterodimer of evolved designer-recombinases precisely excises a human genomic DNA locus. Nucleic Acids Res. (2019) https://doi.org/10.1093/nar/gkz1078.

36. M. H. Hanewich-Hollatz, Z. Chen, L. M. Hochrein, J. Huang, N. A. Pierce, Conditional Guide RNAs: Programmable Conditional Regulation of CRISPR/Cas Function in Bacterial and Mammalian Cells via Dynamic RNA Nanotechnology. ACS Central Science 5, 1241–1249 (2019).

37. M. P. Zeidler, et al., Temperature-sensitive control of protein activity by conditionally splicing inteins. Nat. Biotechnol. 22, 871–876 (2004).

38. N. Dissmeyer, Conditional Modulation of Biological Processes by Low-Temperature Degrons. Methods Mol. Biol. 1669, 407–416 (2017).

39. K. M. Bonger, L.-C. Chen, C. W. Liu, T. J. Wandless, Small-molecule displacement of a cryptic degron causes conditional protein degradation. Nat. Chem. Biol. 7, 531–537 (2011).

40. B. Nabet, et al., The dTAG system for immediate and target-specific protein degradation. Nat. Chem. Biol. 14, 431–441 (2018).

41. M. Schapira, M. F. Calabrese, A. N. Bullock, C. M. Crews, Targeted protein degradation: expanding the toolbox. Nat. Rev. Drug Discov. 18, 949–963 (2019).

42. D. G. Gibson, et al., Enzymatic assembly of DNA molecules up to several hundred kilobases. Nat. Methods 6, 343–345 (2009).

43. J. G. Doench, et al., Optimized sgRNA design to maximize activity and minimize off-target effects of CRISPR-Cas9. Nat. Biotechnol. 34, 184–191 (2016).

44. K. Katoh, D. M. Standley, MAFFT multiple sequence alignment software version 7: improvements in performance and usability. Mol. Biol. Evol. 30, 772–780 (2013).

45. F. Port, H. M. Chen, T. Lee, S. L. Bullock, Optimized CRISPR/Cas tools for efficient germline and somatic genome engineering in Drosophila. Proc. Natl. Acad. Sci. U. S. A. 111, E2967–76 (2014).

46. M. Van Doren, A. L. Williamson, R. Lehmann, Regulation of zygotic gene expression in Drosophila primordial germ cells. Curr. Biol. 8, 243–246 (1998).

47. D. A. Theilmann, S. Stewart, Molecular analysis of the trans-activating IE-2 gene of Orgyia pseudotsugata multicapsid nuclear polyhedrosis virus. Virology 187, 84–96 (1992).

